# MAGMa: Your Comprehensive Tool for Differential Expression Analysis in Mass-Spectrometry Proteomic Data

**DOI:** 10.1101/2024.06.24.600424

**Authors:** Shagun Gupta, Jin Joo Kang, Yu Sun, Yugandhar Kumar, Mateusz Wagner, Will Comstock, James Booth, Marcus Smolka, Haiyuan Yu

## Abstract

Proteomics, the study of proteins and their functions, plays a vital role in understanding biological processes. In this study, we sought to address the challenges in analyzing complex proteomic datasets, where subtle changes in protein abundance are difficult to detect. Utilizing a newly developed tool, **M**aximal **A**ggregation of **G**ood protein signal from **Ma**ss spectrometric data (**MAGMa**), we demonstrated its superior performance in accurately identifying true signals while effectively filtering out noise. Here we show that MAGMa strikes a balance between sensitivity and specificity on benchmarking datasets, offering a robust solution for analyzing various quantitative proteomic datasets. These findings advance the field by providing researchers with a powerful tool to uncover subtle changes in protein abundance, contributing to our understanding of complex biological systems and potentially facilitating the discovery of new therapeutic targets.

## Introduction

Relative quantitative proteomics has been instrumental in driving discovery-based explorations into the dynamic expression of proteins and their modifications across various biological and disease states. In recent years, it has played a pivotal role in elucidating insights across diverse domains, ranging from the pathobiology of SARS-CoV2 viral infection ^1–3^ to the molecular mechanisms underlying RNA polymerase II transcription machinery ^4–6^. Commonly used techniques in the realm of relative quantitative proteomics, performed without prior knowledge of the peptide or protein composition of the sample, include Label-Free Quantification (LFQ), which, as the name implies, enables relative quantitation without relying on any chemical labels. Additionally, techniques such as Tandem Mass Tag (TMT) ^7^ or isobaric Tags for Relative and Absolute Quantitation (iTRAQ) ^8, 9^ facilitate isobaric labeling of peptides post-translation. Another approach, stable isotope labeling in cell culture (SILAC),^10^ involves growing cells with different non-radioactive isotopes in vivo during translation, enabling comparison of proteins or protein expression profiles across different biological states.

In TMT/iTRAQ, quantification occurs post-fragmentation of selected ions, with subsequent scans distinguishing reagents labeled under different conditions based on their molecular mass distribution. However, ratio compression can distort quantification due to interference from co-eluting peptides, often necessitating TMT-based quantification at the third scan stage^11^. Conversely, SILAC-based quantification occurs at the first scan stage, where heavy-labeled and light-labeled peptides appear as distinct peak doublets. LFQ-based quantification also takes place at the first scan stage. However, LFQ quantification faces challenges such as low reproducibility, sparse data with numerous missing values, and biases introduced during sample preparation and machine runs.

TMT mitigates some of these challenges by reducing missing values that may accrue if samples were run separately, thanks to its multiplexing capabilities allowing multiple samples to be tested simultaneously in a single machine run. With 18 isobaric reagents, TMT facilitates comparison of 18 distinct biological states in parallel. In contrast, SILAC offers limited multiplexing capabilities, interpreting up to three distinct biological conditions using light, medium, and heavy isotopic combinations in cell culture media.

Analysis of diverse dataset types such as TMT and SILAC, involves extensive preprocessing and robust statistical methods to accurately identify differentially expressed proteins or peptides. In a single TMT experiment, complex data structures emerge, with multiple identifications linked to peptide sequences and, consequently, proteins. Thus, a crucial decision arises regarding the utilization of uniquely associated identifications, necessitating the exclusion of those shared between proteins. Moreover, reporter ion signals associated with these identifications, that are ultimately used for quantification, can vary significantly, posing challenges in achieving consistent measurements. Determining quality filtering and aggregation strategies for accurate and precise estimation of peptide or protein presence becomes particularly challenging. Additionally, techniques like fractionation^12^, aimed at enhancing coverage, introduce further complexities, as peptide identifications may span consecutive fractions, leading to diverse quantification signals subject to different kinds of biases. Decisions on how to deal with these biases, among others, in the data processing pipeline carry significant consequences, underscoring the complexity and importance of each analytical step. Furthermore, most statistical methods can easily detect large shifts in protein abundance under different conditions, particularly for highly abundant proteins or those with numerous uniquely associated identifications. However, detecting subtle abundance changes from a few measurements poses a greater challenge. Similar challenges persist across SILAC datasets, highlighting the need for meticulous consideration of biases and dataset structures to ensure optimal analysis outcomes.

The pipeline for processing these datasets to identify and calculate relative abundance of peptides and hence proteins include a few major steps. These include tools that match theoretical spectra derived from possible peptide sequences to experimental second stage scans from the mass spectrometer. These are followed up on by tools that prioritize and filter these Peptide to Spectrum Match (PSM’s) and subsequently use the quantification measures associated with them to calculate relative protein expression profiles. On the computational front, there are a wide variety of pipelines available for the end-to-end processing of these datasets. Some popular search tools include COMET^13^ (a direct open source descendant of University of Washington’s academic version of SEQUEST), SEQUEST^HT^ (the commercial version coupled with Proteome Discoverer (PD)^14^ software developed by Thermo Fisher Scientific), MSFragger^15^ (developed by Nesvizhskii lab at University of Michigan) and Andromeda^16^ (coupled with MaxQuant^17^, a popular platform for analysis of many types of proteomic datasets). Several popular tools exist already that are used for differential expression analysis of these datasets that are coupled with some of the search tools mentioned above. For Tandem Mass Tag (TMT)-based quantification, some popular tools include Proteome Discoverer (PD)^14^, MSstatsTMT ^18^ developed by Vitek lab at Northeastern University, TMT-Integrator (coupled with MSFragger), and PSM-based methods ^19, 20^ that utilize all the reporter intensities of all PSMs associated with a protein to quantify it relative to all other proteins in your dataset. For SILAC datasets, all these tools, with the inclusion of MaxQuant and exclusion of MSstatsTMT can be employed for quantification. Apart from these, several publications cite in- house scripts ^21, 22^ used to process these datasets contributing to the expanding choices. With constant evolution of both instrumentation and bioinformatics tools applied for protein identification and quantification over the years, there is no consensus on how to process these datasets in this fragmented space. Although efforts have been made to systematically compare various machines and data acquisition strategies for proteomic datasets, ^18, 23, 24^ aiming to inform analysis strategies, there is currently a lack of a tool that offers flexibility in applying the crucial insights gleaned from these comparisons across a wide range of dataset types.

In this article, we introduce our tool, **M**aximal **A**ggregation of **G**ood protein signal from **Ma**ss spectrometric data (**MAGMa**), which addresses the challenge of detecting small shifts in abundances with few measurements by outperforming popular tools in detecting true signals while effectively identifying noise. MAGMa achieves a balance between sensitivity and specificity upon evaluation with benchmarking datasets, showcasing its robust performance across diverse applications, including the analysis of previously published biologically relevant datasets, where it uncovers pertinent biology that other tools might overlook. It offers a user-friendly web server with customizable statistical assumptions tailored to specific analyses and supports processing of popular quantitative proteomic datasets (TMT and SILAC) (Fig. 1a). Additionally, MAGMa provides downstream visual assessment through features like volcano plots, enabling users to set thresholds and apply filters. The website can be accessed at the URL: https://magma.yulab.org.

**Figure 1:**
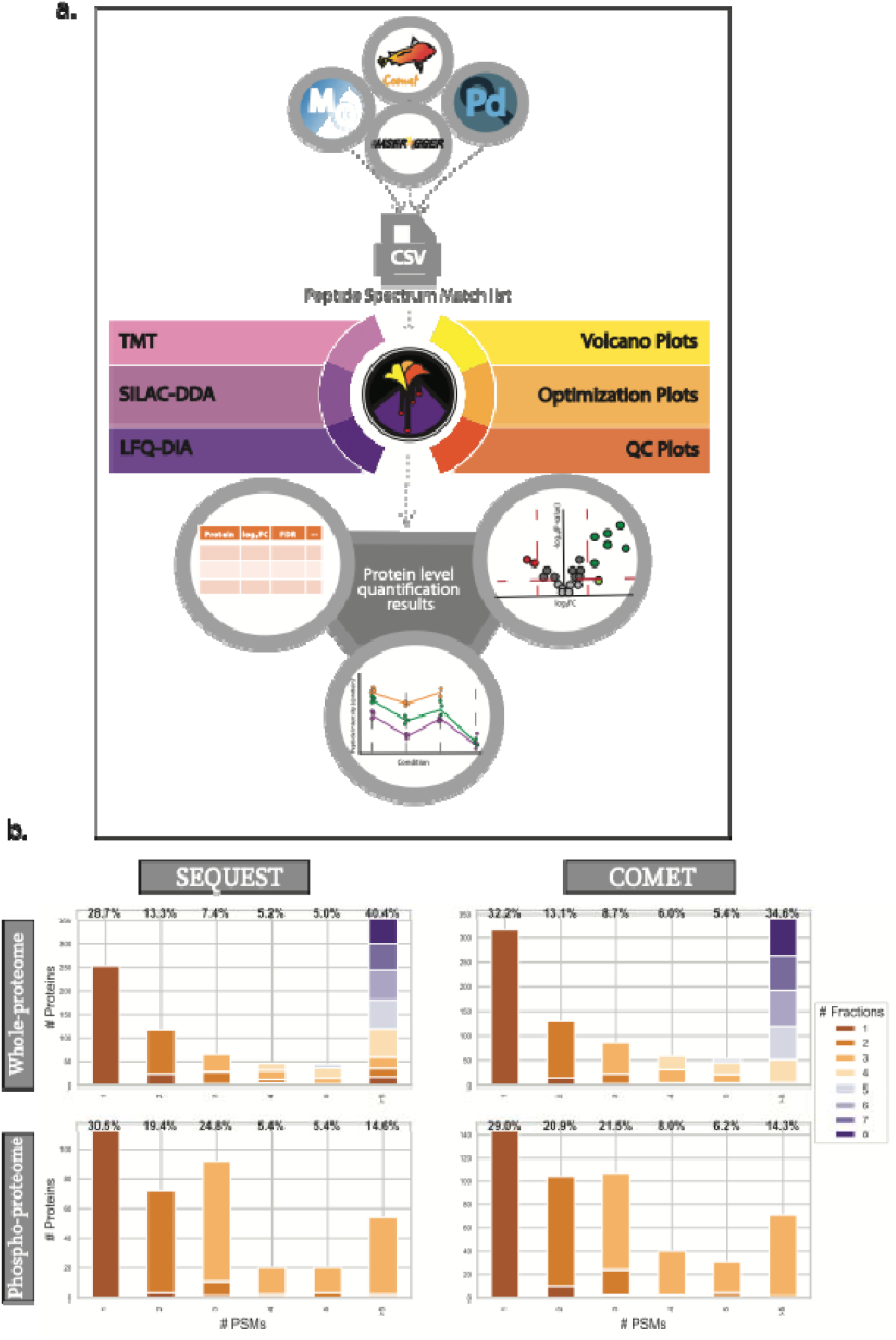
Schematic explanation of functionalities and motivation for the development of MAGMa. (a) Workflow explaining the functionalities available with MAGMa webserver, including the types of datasets (TMT, SILAC, DIA-LFQ) that can be processed using several search tools (SEQUEST, COMET etc.) as well as downstream capabilities with volcano plot-based visualizations and quality control plots. (b) Distribution of proteins identified and quantified in benchmarking datasets across different search tools utilized in the manuscript depicted by the number of peptide-to-spectrum matches (PSMs) and the number of fractions in which these proteins were found, demonstrating the utility of looking at lowly represented proteins.

## Result

### The following section is going to describe the tool MAGMa behind the webserver and the metrics used to evaluate its performance in comparison to existing methods

We employed two types of benchmarking datasets: an in-house generated isobaric labeled (TMT) whole proteome dataset and a published phosphoproteome dataset ^23^, both containing human and yeast whole cell lysates. This deliberate selection aimed to capture diverse challenges encountered in real-world scenarios, particularly in discerning contaminating signals from co-eluting peptides, commonly referred to as ratio compression. Our approach, colloquially termed as a benchmarking dataset, employs the yeast proteome to simulate true signals aimed at detection by our analytical machinery. Conversely, the human proteome models the prevalent background noise that often distorts genuine signals in practical datasets.

While numerous studies assess tool performance using benchmark datasets primarily through calculated fold change (FC), relying solely on this parameter overlooks a crucial aspect: the adjusted P-value or False Discovery Rate (FDR). A substantial observed FC doesn’t necessarily negate the possibility of chance influencing the observation. Hence, it’s imperative to compute the likelihood of observing a difference as significant as the one detected, even if the compared observations originate from populations with identical means. This calculation is further adjusted to accommodate the increased risk of false statistical inferences when running tests on thousands of proteins in parallel.

Therefore, a balanced consideration of both FC and FDR is indispensable for drawing meaningful conclusions. In real experimental scenarios, the evaluation of biological significance typically involves volcano plots, which not only incorporate FC but also assess the likelihood of a protein being differentially expressed in each comparison.

Merely comparing the FC distribution for all yeast proteins isn’t sufficient (Supplementary Fig. 2- 5 a), as it often leads to similar performance outcomes across various methods and search tool combinations. Evaluating performance based solely on FC and FDR thresholds for yeast proteins would also fall short, as it would provide insight into Type I errors but fail to address Type II errors unless simultaneously assessing performance in terms of human proteins modeling the background noise.

Consequently, our evaluation methodology focuses on how effectively the tools identify enriched yeast proteins and unchanged human proteins, both quantifiable from the benchmarking datasets, under the same evaluation criteria.

Across the benchmarking datasets employed in this study, a consistent trend emerged: roughly 50-70% of the dataset comprises proteins with limited representation, evidenced by few peptide-spectrum matches (PSMs) or uniquely assigned peptide sequences, irrespective of the search engine utilized. Notably, this trend is particularly pronounced in phosphoproteome datasets, where proteins with scant identifications predominate (Fig. 1b). Given that most of the analyzed datasets feature low-abundance proteins, the precise and accurate quantification of these entities assumes paramount importance.

Therefore, evaluating MAGMa’s performance using both calculated fold change (FC) and false discovery rate (FDR) is crucial. This evaluation should encompass not only the "easier" scenarios, characterized by highly abundant yeast proteins exhibiting significant changes between biological states but also the more challenging instances involving low-abundance yeast proteins demonstrating borderline significant alterations. Simultaneously, a similar assessment should be conducted on human proteins to gauge MAGMa’s performance comprehensively.

In assessing tool performance with each dataset, three types of comparisons are possible. When comparing conditions, A and B for yeast proteins, the anticipated fold change (FC) is 2.5; for conditions B and C, the expected FC is 4; and for conditions A and C, the anticipated FC is 10. Similarly, for human proteins, the expected FC remains at 1 across all condition comparisons (Fig. 2a, 3a). True positive, false positive, true negative, and false negative numbers are calculated using yeast and human proteins that qualify as differentially enriched by employing commonly accepted thresholds of FC at 2 and a False Discovery Rate (FDR) of 5%. These numbers are then used to calculate subsequent scores like F1 and G-mean (see Methods) to evaluate tool performance for each condition comparison. For the comparison involving a ten-fold versus four-fold spike-in of yeast aliquot, we adjust the FC threshold to 1.5 to capture changes close to a FC of 2.5. The performance evaluation encompasses MAGMa alongside several open-source and published tools, executed across various search engines to substantiate subsequent claims.

**Figure 2:**
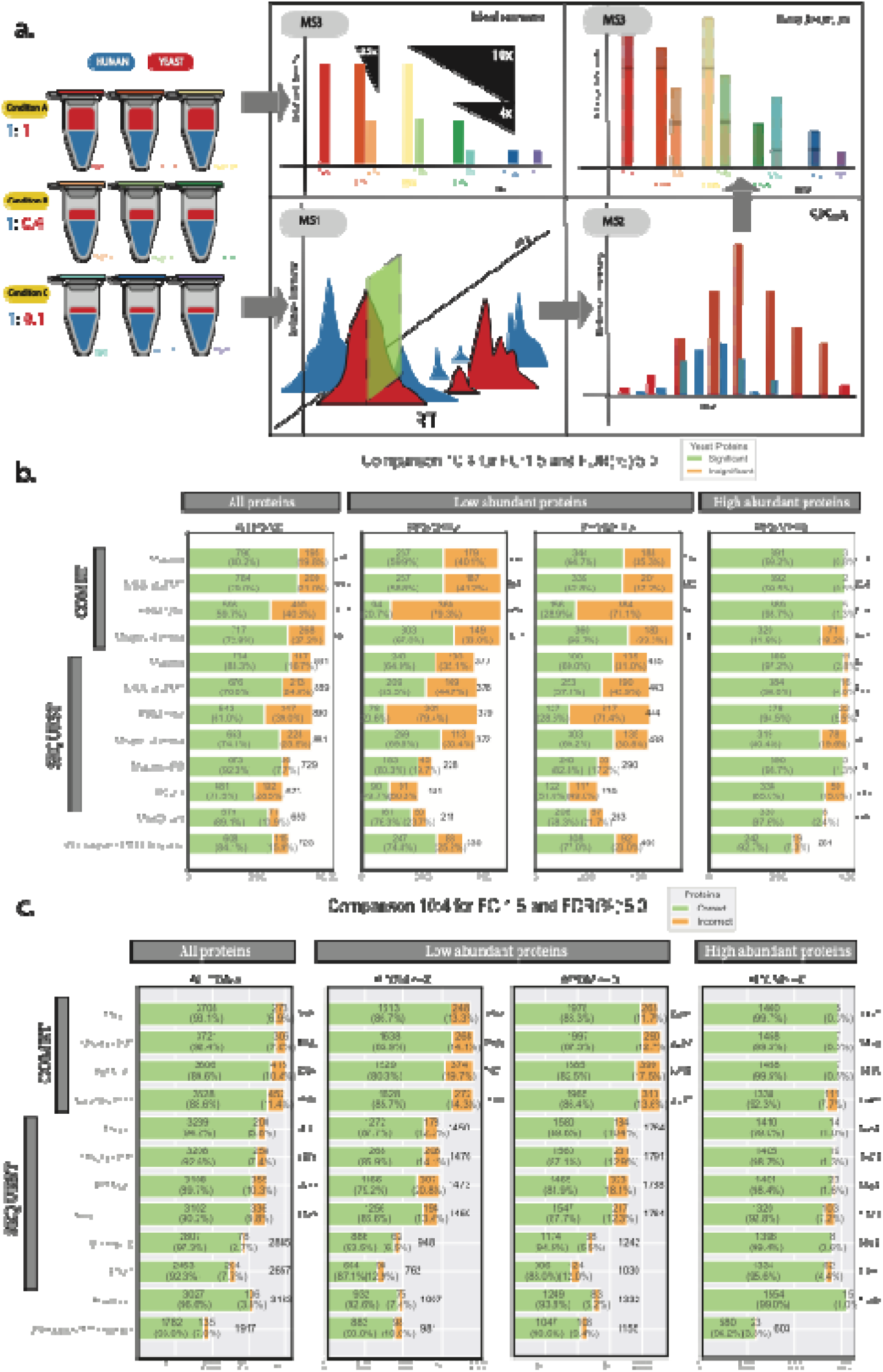
Performance evaluation of open-source and proprietary software(s) with MAGMa on the in-house benchmarking dataset. (a) Description of the TMT setup used for generating the benchmarking dataset. It includes an explanation of how ratio compression is tested within this setup, highlighting the potential impact on quantification accuracy. (b) Performance evaluation of various tools specifically on yeast proteins. It focuses on the tools’ ability to detect small shifts in protein abundance, with an expected fold change (FC) of 2.5, comparing results across both low and highly abundant proteins. For each bar, in green are the number and proportion of yeast proteins that pass FC: 1.5 and FDR: 5% thresholds and in orange are number and proportion of yeast proteins that do not pass these thresholds. The number at the top of each bar is the total number of yeast proteins. (c) Performance evaluation of tools on all proteins (yeast and human). For each bar, in green are the number and proportion of yeast proteins that pass FC: 1.5 and FDR: 5% thresholds and human proteins that do not pass these thresholds (correct calls). In orange are number and proportion of yeast proteins that do not pass these thresholds and number of human proteins that pass these thresholds (incorrect calls). The number at the top of each bar is the total number of proteins.

### MAGMa allows enhanced detection of small abundance shifts in whole proteome TMT benchmark datasets

In the benchmarking datasets discussed in this manuscript, we evaluate subtle abundance shifts by comparing spike-in conditions with an expected fold change (FC) of 2.5. Assessing tool performance based on the number and proportion of yeast and human proteins meeting FC thresholds of 1.5 and a False Discovery Rate (FDR) of 5%, MAGMa outperforms conventional tools (Fig. 2-3, b-c). Furthermore, when considering metrics such as F1 score, reflecting the balance between precision and recall, and G-mean, which evaluates the balance of sensitivity and specificity, MAGMa consistently surpasses other tools (Table 1-2). This superiority holds true across various search tools, evident in both the in-house constructed (Fig. 2b-c, Table 1) and published datasets (Fig. 3b-c, Table 2).

**Table 1:** Comparison analysis on in-house benchmarking dataset on the ability to detect very small shifts in protein quantifications. Cutoffs of FC:1.5, FDR: 5% were used for this comparison where yeast proteins have expected FC of 2.5, and human proteins have expected FC of 1.

|  |  | All proteins |  |  |  | Low abundant proteins |  |  |  |  |  |  |  | High abundant proteins |  |  |  |
| --- | --- | --- | --- | --- | --- | --- | --- | --- | --- | --- | --- | --- | --- | --- | --- | --- | --- |
| Tool |  | All PSMs |  |  |  | <=2 PSMs |  |  |  | <=3 PSMs |  |  |  | >=5 PSMs |  |  |  |
|  |  | G-mean | Recall | F1 | Accuracy | G-mean | Recall | F1 | Accuracy | G-mean | Recall | F1 | Accuracy | G-mean | Recall | F1 | Accuracy |
| COMET | Magma | **0.884** | **0.802** | **0.853** | **0.931** | **0.755** | **0.599** | **0.683** | **0.867** | **0.786** | **0.647** | **0.723** | **0.883** | **0.995** | 0.992 | **0.994** | **0.997** |
|  | MSstatsTMT | 0.874 | 0.790 | 0.837 | 0.924 | 0.745 | 0.588 | 0.666 | 0.859 | 0.772 | 0.628 | 0.700 | 0.873 | 0.995 | **0.995** | 0.991 | 0.995 |
|  | PSM-ratio | 0.770 | 0.597 | 0.739 | 0.896 | 0.453 | 0.207 | 0.335 | 0.803 | 0.535 | 0.289 | 0.439 | 0.825 | 0.993 | 0.987 | 0.991 | 0.995 |
|  | Magma-Limma | 0.771 | 0.607 | 0.726 | 0.886 | 0.716 | 0.535 | 0.640 | 0.857 | 0.723 | 0.543 | 0.654 | 0.864 | 0.847 | 0.719 | 0.835 | 0.923 |
| SEQUEST | Magma | **0.903** | **0.833** | **0.880** | **0.942** | **0.787** | **0.649** | **0.729** | **0.877** | **0.815** | **0.690** | **0.765** | **0.896** | **0.985** | **0.973** | **0.982** | **0.990** |
|  | MSstatsTMT | 0.864 | 0.760 | 0.840 | 0.926 | 0.730 | 0.553 | 0.668 | 0.859 | 0.744 | 0.571 | 0.687 | 0.871 | 0.978 | 0.960 | 0.976 | 0.987 |
|  | PSM-ratio | 0.780 | 0.610 | 0.754 | 0.897 | 0.452 | 0.206 | 0.337 | 0.792 | 0.534 | 0.286 | 0.440 | 0.819 | 0.972 | 0.945 | 0.970 | 0.984 |
|  | Magma-Limma | 0.813 | 0.673 | 0.779 | 0.902 | 0.757 | 0.597 | 0.695 | 0.866 | 0.764 | 0.605 | 0.708 | 0.877 | 0.864 | 0.748 | 0.853 | 0.928 |
|  | Magma-PD | **0.956** | **0.923** | **0.945** | **0.973** | **0.885** | **0.803** | **0.855** | **0.935** | **0.901** | **0.828** | **0.876** | **0.945** | **0.992** | **0.987** | **0.990** | **0.994** |
|  | PD 2.3 | 0.843 | 0.715 | 0.825 | 0.923 | 0.701 | 0.497 | 0.647 | 0.871 | 0.711 | 0.510 | 0.663 | 0.880 | 0.921 | 0.850 | 0.915 | 0.956 |
| MaxQuant |  | 0.937 | 0.891 | 0.916 | 0.966 | 0.860 | 0.763 | 0.811 | 0.926 | 0.874 | 0.783 | 0.832 | 0.938 | 0.985 | 0.976 | 0.978 | 0.990 |
| MSFragger TMTIntegrator |  | 0.909 | 0.841 | 0.900 | 0.930 | 0.855 | 0.748 | 0.834 | 0.900 | 0.868 | 0.770 | 0.851 | 0.906 | 0.957 | 0.927 | 0.955 | 0.962 |

**Table 2:**
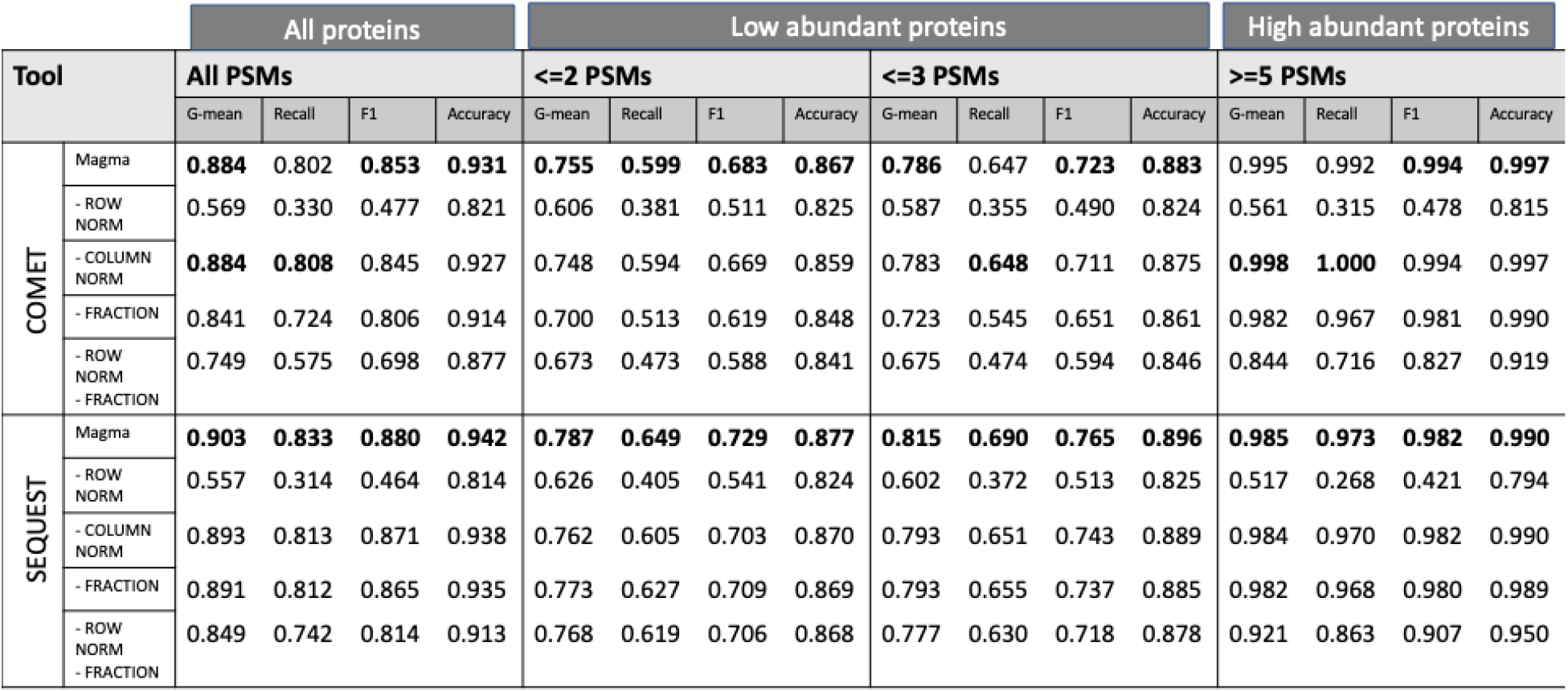
Comparison analysis on published phospho-proteome benchmarking dataset on the ability to detect very small shifts in protein quantifications. Cutoffs of FC:1.5, FDR: 5% were used for this comparison where yeast proteins have expected FC of 2.5, and human proteins have expected FC of 1.

**Figure 3:**
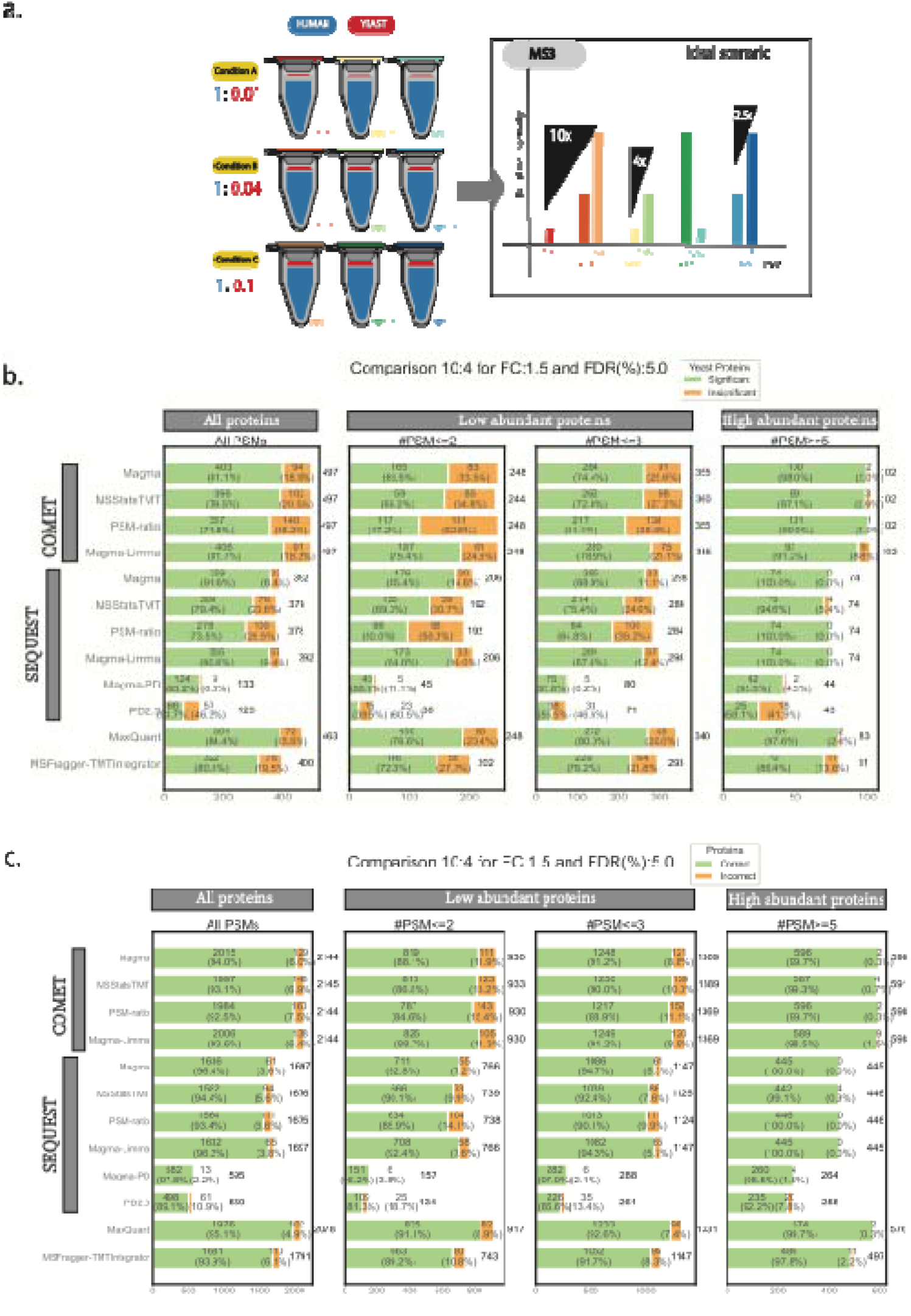
Performance evaluation of open-source and proprietary software(s) with MAGMa on published phosphoproteome benchmarking dataset. (a) Description of the TMT setup used for generating the benchmarking dataset. (b) Performance evaluation of various tools specifically on yeast proteins. It focuses on the tools’ ability to detect small shifts in protein abundance, with an expected fold change (FC) of 2.5, comparing results across both low and highly abundant proteins. For each bar, in green are the number and proportion of yeast proteins that pass FC: 1.5 and FDR: 5% thresholds and in orange are number and proportion of yeast proteins that do not pass these thresholds. The number at the top of each bar is the total number of yeast proteins. (c) Performance evaluation of tools on all proteins (yeast and human). For each bar, in green are the number and proportion of yeast proteins that pass FC: 1.5 and FDR: 5% thresholds and human proteins that do not pass these thresholds (correct calls). In orange are number and proportion of yeast proteins that do not pass these thresholds and number of human proteins that pass these thresholds (incorrect calls). The number at the top of each bar is the total number of proteins.

Prominently, MAGMa excels in accurately and precisely detecting significant FC shifts across other condition comparisons and upon employing more stringent FC and FDR thresholds per comparison (Supplementary Fig. 1-4, b-c, Supplementary Table 1-4).

### MAGMa displays superior performance for enhanced quantification of low-abundance proteins

In this manuscript, low-abundance proteins are characterized as those with three or fewer unique identifications or Peptide Spectrum Matches (PSMs) detectable by the mass spectrometer. Evaluating MAGMa’s performance on these proteins across both in-house and publicly available benchmarking datasets, employing various search strategies, reveals its consistent superiority over other tools. This enhanced performance persists across all three condition comparisons possible with the benchmarking datasets (Fig. 2-3, b-c, Table 1-2, Supplementary Fig. 1-4, b-c, Supplementary Table 1-4).

A direct comparison of MAGMa, PSM-based methods, and MSstatsTMT on proteins exhibiting not only low abundance but also minimal detectable shifts underscores MAGMa’s efficacy. MAGMa identifies more yeast proteins as significant in such comparisons without erroneously classifying human proteins as significant under the same FC and FDR thresholds. This superiority is further underscored by metrics such as F1 and G-mean (Table 1-2). Upon investigating the underlying factors driving this performance boost in terms of FC, FDR, or both, we observe that FDR predominantly influences the trend, as methods exhibit similar performance in terms of FC alone (Supplementary Fig. 5a).

### The synergistic impact of row normalization and fractional value retention in driving MAGMa’s superior performance

Using the metrics outlined in the preceding section, we delve into the specific strategies that underpin MAGMa’s enhanced performance. Our analysis reveals that this improvement stems from a combination of subtle yet pivotal alterations in how we compile and analyze the dataset for differential expression analysis.

The composition of TMT datasets, whether derived from real biological samples or artificially constructed like the benchmark datasets utilized in this study, is complex. A protein’s quantification may stem from multiple peptides uniquely assigned to it, with the possibility of detection occurring multiple times within the same machine run. This complexity is further compounded when considering fractionation, as peptides can be identified and quantified across various machine runs or fractions. Within the subset of the benchmarking dataset comprising low-abundance proteins, approximately 40-64% exhibit representation across multiple fractions. Notably, these peptides may vary significantly in terms of associated quantification values per TMT tag. For instance, a single protein (e.g., Yeast Uniprot ID: P10592) may have one peptide reported at a scale of 2^10, while another peptide may register at 2^20 (Fig. 4b-c).

**Figure 4:**
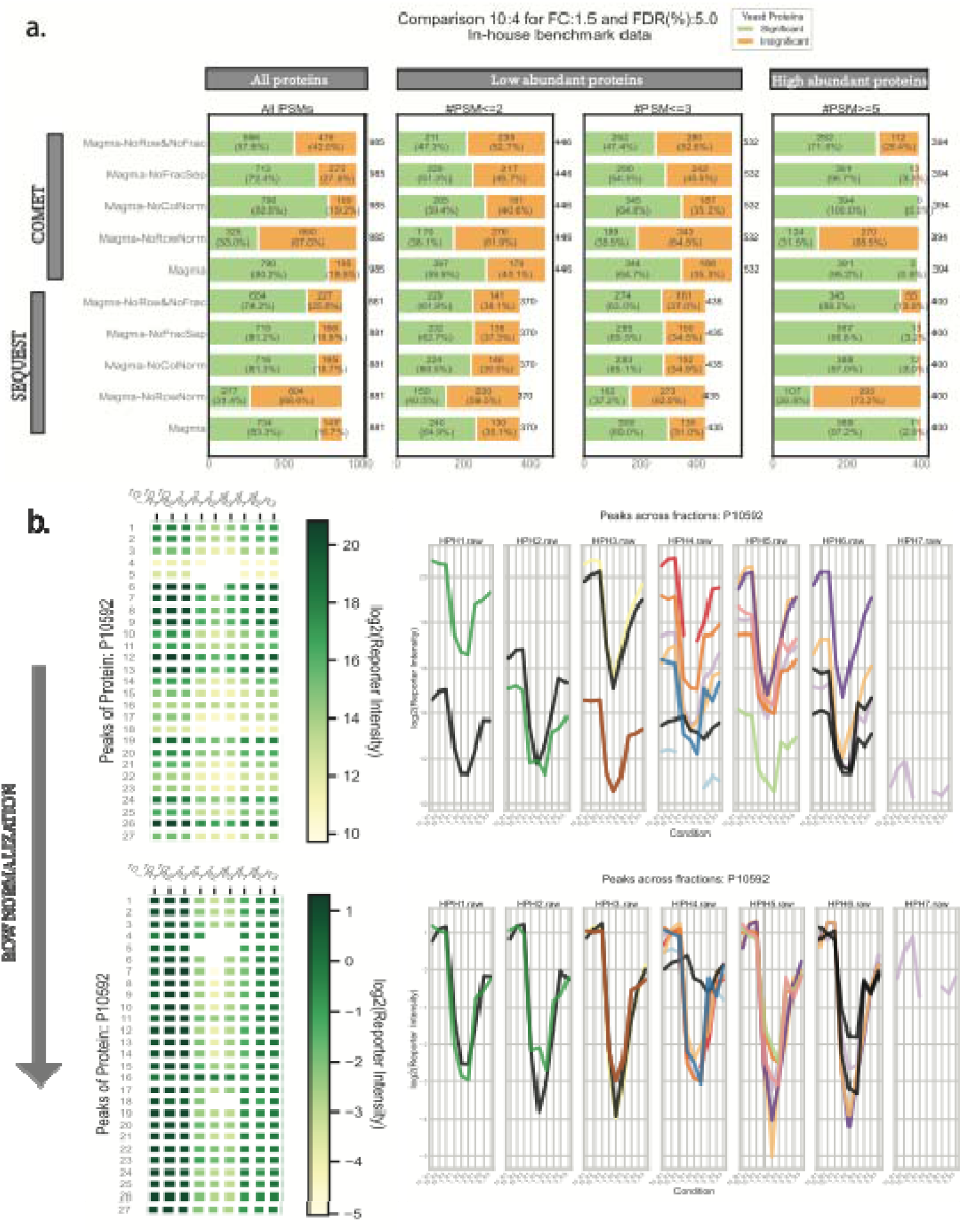
Ablation study highlighting key drivers of MAGMa’s performance. (a) Examination of how different MAGMa tool specifications influence performance. The effects on overall performance metrics are highlighted to identify the key drivers behind MAGMa’s effectiveness. For each bar, in green are the number and proportion of yeast proteins that pass FC: 1.5 and FDR: 5% thresholds and in orange are number and proportion of yeast proteins that do not pass these thresholds. The number at the top of each bar is the total number of yeast proteins. (b) Demonstration of the impact of row normalization on the consistency of measurements associated peaks, which are defined by the unique combination of peptide sequence, charge, and modifications, across different fractions.

A key challenge when summarizing multiple data points, such as peptide quantifications, to the protein level is the risk of overlooking individual peptide characteristics (row-specific features). However, an often-overlooked insight is that ratios between conditions tagged with different TMT tags tend to remain consistent for a protein, despite potential variations in the scale of individual peptide measurements. This consistency is even more pronounced for peptide measurements across different fractions. Referred to as row normalization, this process entails dividing each row of the dataset—following quality-based filtering (refer to Methods)—by the row’s average to preserve peptide-specific characteristics.

MAGMa adopts a strategy of summarizing the dataset to the protein level per fraction. This approach enables the comprehensive utilization of increased coverage resulting from fractionation in subsequent significance assessments (refer to Methods).

We investigate the synergistic effect of row normalization and the retention of fraction-separated values on MAGMa’s performance, particularly in challenging scenarios. Through an ablation study, we assess MAGMa’s performance using the same metrics outlined earlier in the manuscript. Our findings highlight that MAGMa’s performance is influenced by various factors, but the most significant impact occurs when fraction-separated values are retained without row normalization. This impact is particularly pronounced in detecting subtle shifts in abundance (Fig. 4a, Table 3). Importantly, this trend persists across both benchmarking datasets evaluated using different search tools.

**Table 3:**
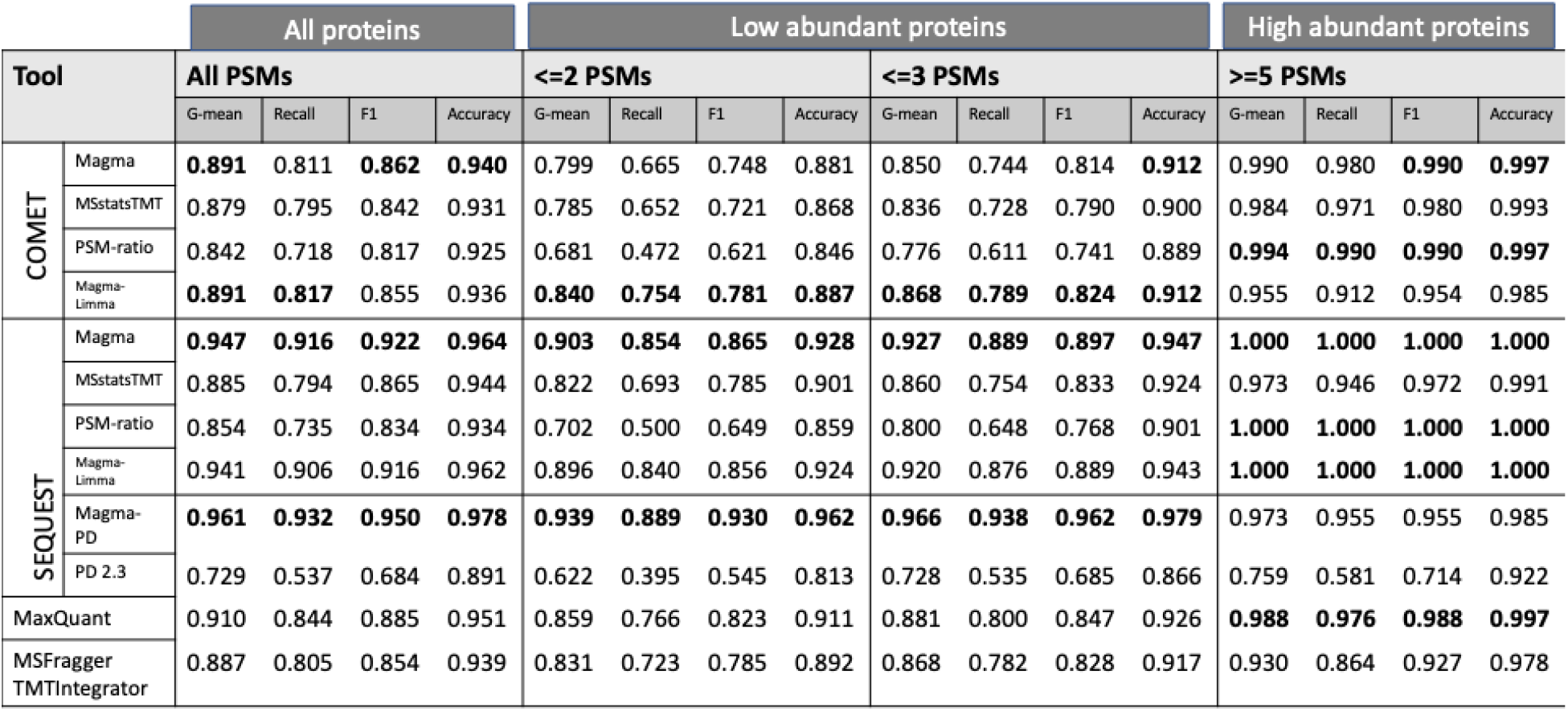
Ablation study on MAGMa on in-house benchmarking dataset on its ability to detect very small shifts in protein quantifications. Cutoffs of FC:1.5, FDR: 5% were used for this comparison where yeast proteins have expected FC of 2.5, and human proteins have expected FC of 1.

Interestingly, the discernable impact observed with fraction-separated values and without row normalization is not replicated when summarizing identifications across fractions in a manner akin to MSstatsTMT while still employing row normalization. Furthermore, the performance of MAGMa with no fraction-based separation combined with no row normalization mirrors that of fraction separation combined with no row normalization. This observation challenges the notion that MAGMa’s performance improvement is solely driven by a larger sample size per low-abundance protein, leading us to conclude that row normalization and the retention of a larger dataset by treating peptide identifications across fractions as independent are vital when summarizing and subsequently conducting differential expression analysis.

### MAGMa web server allows users to choose the best analysis strategies across several data acquisition strategy types

The webserver MAGMa allows users to choose from several ways for analyzing their TMT, or SILAC datasets. Users can choose to employ the insights derived from the comparative analyses discussed above on a wide variety of TMT and SILAC datasets ranging from immunoprecipitation (IP) experiments to phosphoproteomic experiments which necessitate peptide and phosphorylation-site level aggregation of information.

In various experimental setups, such as comparisons between drug treatments or mutations and their wild-type counterparts in IP experiments, conducting parallel processing of multiple condition comparisons is advantageous. However, the ability to dynamically visualize such comparisons simultaneously is lacking in existing tools to our knowledge. The MAGMa web server uniquely offers this functionality, enabling users to assess the impact of different thresholds by visualizing the effects of multiple filters (such as fold change, false discovery rate, and number of peptide-spectrum matches) and protein/peptide-level statistics across all comparisons ^25^. This capability is exemplified through the application of MAGMa in a published chemoproteomic workflow utilizing TMT to pinpoint precise protein binding sites of photoaffinity probes in cells. Here, the comparison of two conditions (enrichment of different photoaffinity probes compared to a control) informs the third comparison (differential enrichment of one affinity probe versus another), with results visualized through volcano plots. (Fig 5).

**Figure 5:**
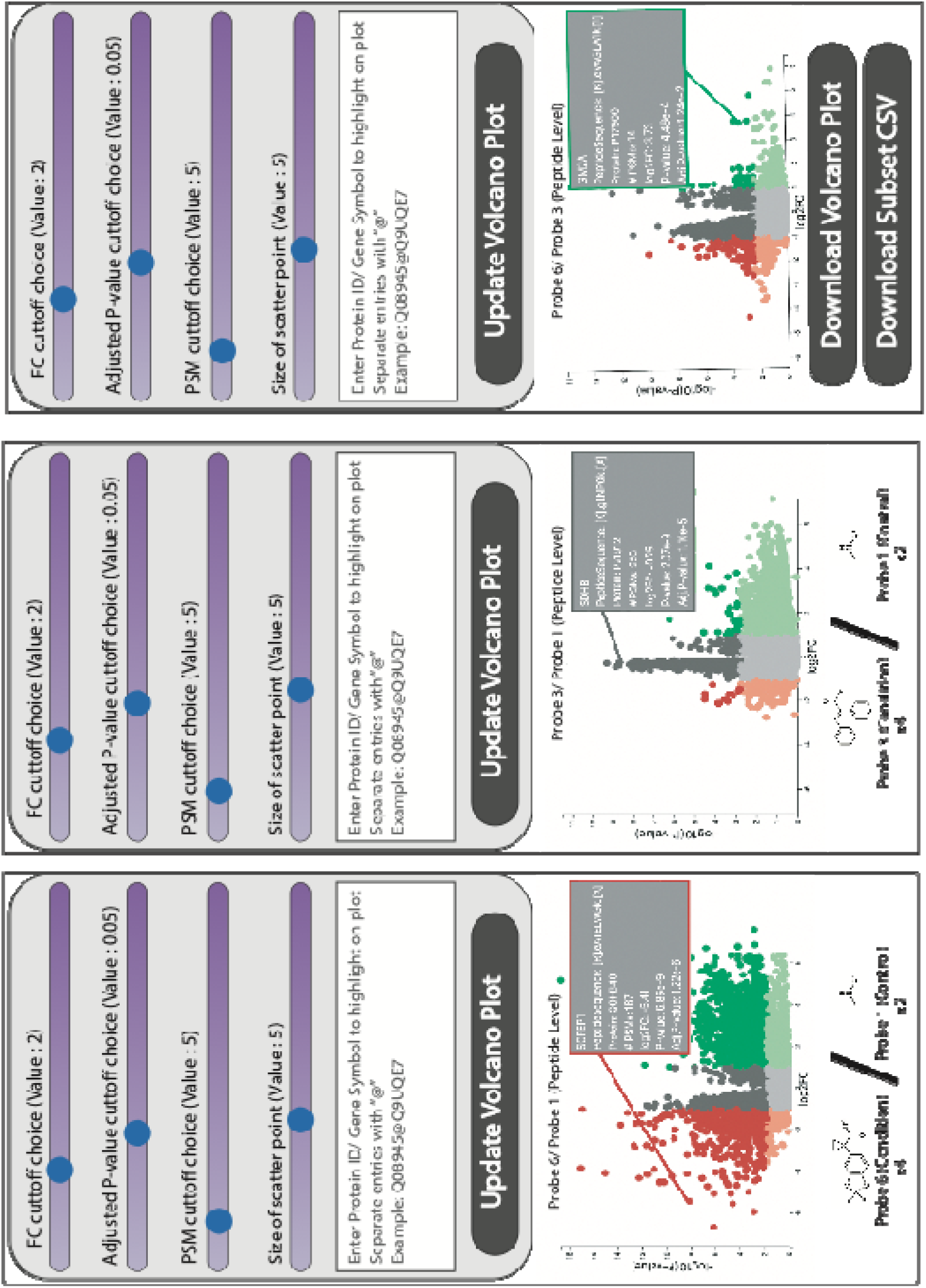
Dynamic visualization of interdependent condition comparisons with MAGMa. Display of MAGMa’s ability to analyze published dataset, requiring parallel comparison of conditions of Probe 3, Probe 6 and Probe 1 (Control), and real time visualization of the impact of commonly used thresholds like FC, FDR, # PSMSs on the relevant volcano plots. For example, the impact of FC associated with Probe 6/ Probe 1 on its volcano plot as well as on Probe 6/ Probe 3 volcano plot.

### MAGMa reveals nuanced biological findings overlooked by other analysis tools

The application of MAGMa to a published dataset comparing tumor samples from Hepatocellular carcinoma (HCC) to peri-tumor samples reveals biological insights consistent with known characteristics of HCC (Fig 6a)^26^. Dysregulation of the PI3K/Akt/mTOR pathway, a hallmark of HCC ^27, 28^, promotes cell growth, survival, and metabolism, with genetic alterations like mutations in PI3K or loss of PTEN expression contributing to pathway activation. Using thresholds of FC>1.5 and FDR<5%, MAGMa identifies enrichment of pathway-associated proteins in peri-tumor samples. Moreover, dysregulation of metabolic pathways, including glycolysis, oxidative phosphorylation, fatty acid metabolism, and amino acid metabolism, is commonly observed in HCC ^27, 29^. This is recapitulated by the enrichment of several proteins (FC<0.67, FDR<5%) belonging to these metabolic pathways in Tumor samples. However, comparison with results from the open-source tool MSstatsTMT highlights that MAGMa captures trends missed by MSstatsTMT, even when using commonly used thresholds of raw P-values (instead of FDR) (Fig 6b-c). This underscores the unique ability of MAGMa to uncover nuanced biological insights that may be overlooked by other analysis tools.

**Figure 6:**
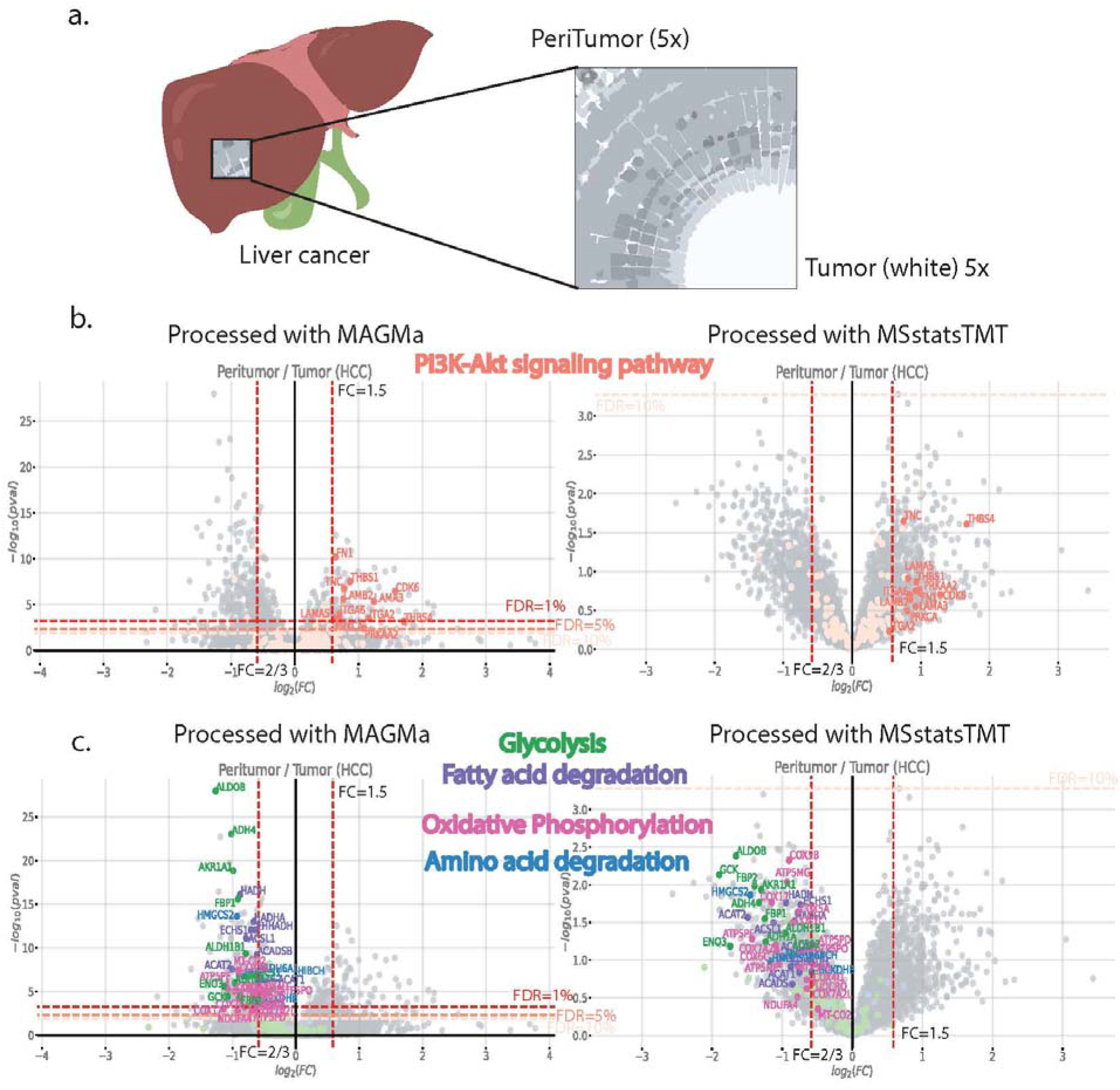
MAGMa’s ability to capture biological trends missed by other tools. (a) Schematic explaining the setup of the published dataset, comparing the peritumor and tumor sections (5 replicates each using TMT10) of formalin fixed paraffin embedded (FFPE) preserved tissue taken from Hepatocellular carcinoma (HCC) specimen (b) Comparison of differential expression analysis done using MAGMa and MSstatsTMT in their ability to capture proteins associated with P3I3K-Akt signaling pathway (red, KEGG: hsa04151). On the volcano plot, analysis is done comparing peritumor (enriched on the right) versus tumor (enriched on the left) at thresholds of FC 1.5 (red vertical dashed lines) and FDR:1% (dark red dashed horizontal line), FDR: 5% (red dashed horizontal line) and FDR: 10% (light red dashed horizontal line). Dark red scatter dots are also labeled and refer to proteins belonging to the pathway that pass the thresholds of FC:1.5, FDR: 5%. Light red scatter dots refer to all other proteins belonging to this pathway that were identified in this experiment. Most of these colored scatter points do not pass raw p-value threshold of 0.05 for MSstatsTMT. (c) Same analysis is done as (b) but using pathways Glycolysis (green, KEGG: hsa00010), Fatty acid degradation (purple, KEGG: hsa00071), Oxidative Phosphorylation (maroon, KEGG: hsa00190), Amino acid degradation (blue, KEGG: hsa00280).

## Discussion

The landscape of tools for relative quantification of analytes is fragmented, spanning from search engine choices for converting mass spectrometry-acquired spectra to post-processing tools aggregating information to the peptide or protein level. However, many existing tools falter when assessing low-abundance targets with few identifications for quantification. Demonstrating superior performance on both low-abundance proteins and those with limited identifications, MAGMa shines when evaluated on TMT benchmarking datasets across multiple search engines. As we consider the future, the inclusion of DIA-LFQ as an additional arm in MAGMa’s website becomes imperative. DIA-LFQ’s cost-effectiveness and competitiveness as a technique, along with shared nuances in data structure, make it a natural fit for MAGMa’s capabilities.

Compared to existing methodologies, MAGMa excels in processing various search types and quantitative proteomic techniques. Key strategies driving this performance boost include row normalization and treating signal spread across fractions as independent contributors to protein abundance. These nuanced yet impactful changes streamline statistical processing, obviating the need for complex models while ensuring maximal proteome coverage.

Furthermore, when applied to a published TMT phosphoproteomic benchmarking dataset, MAGMa demonstrates its ability to outperform other tools, particularly in evaluating phosphorylated proteins. While MAGMa may curate a smaller number of proteins for performance comparisons, it does so by excluding those lacking adequate reporter ion abundances for evaluation. Leveraging empirical Bayes moderation to borrow strength for surrogate variance calculation further enhances performance, although caution is advised due to increased erroneous assessments (Supplementary Fig. 8, supplementary Table 7).

MAGMa’s optimal performance is realized when TMT experimental setups are confined to a single machine run and a single batch of multiplexing labels. However, a limitation lies in its current inability to address batch effects, particularly in setups requiring layouts separated across runs. Future efforts will focus on incorporating model flexibility to address batch effects effectively.

## Method

### Construction of in-house benchmark dataset

#### Yeast (haploid yeast BY4741) sample preparation

The preparation of yeast extract has been previously described in detail (Cell, Tommy V Vo et al., 2016). Strains (BY4741) were grown in YPD (Yeast Extract-Peptone Dextrose) media with appropriate supplements (adenine and glucose) at 30°C overnight. Then, cells were harvested by centrifuge at 700g for 5min. Cells were lysed by using cold glass beads in lysis buffer (50mM tris-HCl pH 7.5, 0.2% tergitol, 150mM NaCl, 5mM EDTA) containing protease inhibitor cocktail (Roche) and 5mM PMSF. Whole cell lysates were collected and normalized by using Bradford assay.

#### Human 293T cell sample preparation

HEK293T cells were grown in DMEM media supplemented 10% Fetal Bovine Serum. Cells were collected and washed 3 times in 10ml DPBS (VWR, 14190144), resuspended in 500ul of NP-40 lysis buffer (50mM HEPES pH 7.5, 150mM NaCl, 5mM EDTA, 1.0% NP-40) and incubated on the ice for 30 min. Cell lysate was sonicated on a sonifier cell disruptor (BRANSON, 500-220-180) for 120 sec at 40% amplitude. Cell extracts were cleared by centrifugation at 16,100g at 4°C for 15 min. whole cell extracts were normalized by Bradford assay. Three different human versus yeast cell extract ratios of 1:1, 1:0.4 and 1:0.1 were prepared by concentration of both cells.

#### TMT sample preparation

TMT procedures were carried out as previously described^3^. Human and yeast mixtures were reduced for 1h at 55°C using 200 mM TCEP. Then, 375 mM iodoacetamide was used to conduct alkylation for 30 minutes at room temperature in the dark. Trypsin Gold, mass spectrometry grade (catalog no. V5280; Promega), was then used to digest the samples at a 1:100 enzyme-to-substrate ratio. The samples were incubated at 37°C for overnight. After that, the Pierce Quantitative Colorimetric Peptide Assay (catalog no. 23275; Thermo Scientific) was used to measure the peptide concentrations. Triethylammonium bicarbonate (TEAB, 1M, catalog no. 90114; Thermo Scientific) was used to resuspend and standardize samples for TMT experiment. Using the TMT10plex Isobaric Mass Tagging Kit (catalog no. 90113; Thermo Scientific), samples were tagged for 1h at room temperature with a label-to-peptide ratio of 20:1. 5% hydroxylamine was added to the labeling reactions to quench them for 15 minutes. The reactions were then pooled and dried using a SpeedVac. The Pierce High pH Reversed-Phase

Peptide Fractionation Kit (catalog no. 84868; Thermo Scientific) was utilized to enrich and fractionate labeled peptides in compliance with the manufacturer’s procedure.

#### LC-MS/MS Measurements

Tryptic peptides were analyzed on Orbitrap Fusion Lumos Tribrid Mass Spectrometer (catalog no. IQLAAEGAAPFADBMBHQ; Thermo Scientific) coupled to an in-house 3 μm C18 resin- (Michrom BioResources) packed capillary column (125 μm × 25 cm), the EASY-nLC 1200 System (catalog no. LC140; Thermo Scientific) was used to analyze fractions. For peptide separation, the following elution gradient and mobile phase were used: Buffer A: 0.1% formic acid in water; buffer B: 0.1% formic acid in 80% acetonitrile; flow rate adjusted to 300 nl min−1. 0–5 min, 5%–8% B; 5–65 min, 8–45% B; 65–66 min, 45%–95% B; 66–67 min, 95% B; 67-68 min, 95%-2% B; 68-72 min 2%-95% B; 72-80 min 95%-5% B. MS^1^ precursors were found at resolution of 120,000 and m/z of 375–1500. MS^n^ data was acquired using the CID-MS2-HCD- MS3 method. For MS^2^ analysis with resolution = 50,000, isolation width = 0.7 m/z, maximum injection duration = 50 ms, and CID collision energy at 35%, precursor ions with charge of 2+ to 7+ were selected. For MS^3^ analysis, six SPS precursors were selected, and ions were fragmented up using 65% HCD collision energy. Spectra was acquired using Tune program v.3.4 (Thermo Scientific) and Thermo Xcalibur Software v.4.4 (catalog no. OPTON-30965; Thermo Scientific).

### Database Searching

#### SEQUEST^HT^

Searches were performed using “.raw” files using Proteome Discoverer (version 2.3.0) with its built-in search engine, SEQUEST^HT^. Tandem mass spectra were searched against the human reference proteome (UP000005640, downloaded on February 10, 2022) and the yeast reference proteome (UP000002311, downloaded on February 10, 2022). Precursor mass tolerance was set to 10 ppm, and fragment mass tolerance to 0.6 Da. For the analysis done in this manuscript, TMT10 was specified as a fixed label on lysine and peptide N termini. For all searches, carbamidomethylation cysteine was set as a fixed modification and oxidation of methionine and N-terminal protein acetylation as variable modifications. Trypsin/P was specified as the proteolytic enzyme with up to two missed cleavage sites allowed and up to three dynamic modifications were allowed. The search was run with following changes in default processing and consensus workflows for TMT-SPS MS3 quantification provided with Proteome Discoverer. The parameter filtering out S/N values under 1.5 (in processing workflow in node Spectrum Selector under Peak Filters) was set to 0. The search was run twice by exporting reporter ion intensities as signal to noise (S/N) values and as raw relative intensities (in consensus workflow in node Reporter Ions Quantifier under Reporter Quantification -> Reporter Abundance Based On). There was a subsequent adjustment to 1% FDR with Percolator to get a PSM list employing a target–decoy approach. The results were exported to a tab delimited text file, ensuring the columns “Sequence” and “Intensity” were included before export for processing with MAGMa. Peptide to protein summarization and subsequent FC, P-value and FDR calculation was performed as implemented by Proteome Discoverer for performance comparison with MAGMa.

The search was performed in a similar manner for the published phosphorylation proteome-wide benchmarking dataset except for searching with phosphorylation of serine, threonine and tyrosine as a variable modification as well as search parameters set according to associated publication accounting for MS3 spectrum acquisition in orbitrap^23^.

#### COMET

Searches were performed after conversion of “.raw” files to mzXML format using RawConverter^30^ (version 1.2.0.1) to account for MS3 level quantification for all the benchmarking dataset used herein. Files were processed with COMET (version 2023.01 rev.0) coupled with Trans Proteomic Pipeline (version 6.2.0 Nacreous), using PeptideProphet to filter at 1% FDR (0.8516 probability for whole proteome in-house and 0.8975 for published phosphoproteome benchmarking dataset). The same reference proteome files were used as mentioned above and run parameters were set to replicate SEQUEST search for fair comparison. These were processed with the -PREC flag in PeptideProphet to get precursor intensity measurements per PSM. Isobaric quantification (per TMT channel) was done using Libra, setting quantification at MS3 with SPS data (Thermo MS3), centroiding with intensity weighted mean. The subsequent files were exported and converted to tab delimited text files for processing with MAGMa.

#### MSFragger

Searches were performed after conversion of “.raw” files to mzML format using RawConverter^30^ (version 1.2.0.1) with MSFragger (version 3.7) coupled with FragPipe (version 19.1) with parameters set to replicate COMET searches for both benchmarking datasets. Isobaric quantification (per TMT channel) was done using TMT-Integrator and Philosopher (version 4.8.1). The files were processed with *Limma*^31^ to get the raw *P*-values and FDR (Benjamini Hochberg correction) per condition comparison.

#### MaxQuant

Searches were performed with “.raw” files using MaxQuant (version 2.3.1.0) and subsequent isobaric quantification (per TMT channel) was done using Perseus (version 2.0.9.0) to get FC, raw *P*-values and FDR (Benjamini Hochberg correction) per condition comparison.

#### Performance Evaluation Metrics

F1-score is the harmonic mean of Precision and Recall, providing a single score that balances false positives and false negatives concerns. This metric is especially useful when dealing with an imbalanced dataset (as is the case with benchmarking dataset) where the number of positive instances (with yeast proteome) are not equal to the number of negative instances (with human proteome).

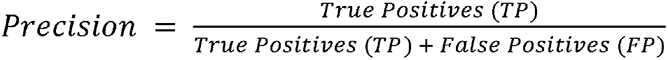

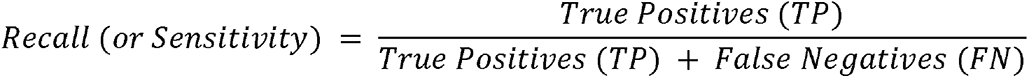

Where,

TP: Number of Yeast proteins that pass the significance threshold of FC and FDR. FP: Number of Human proteins that pass the significance threshold of FC and FDR.

TN: Number of Human proteins that do not pass the significance threshold of FC and FDR.

FN: Number of Yeast proteins that do not pass the significance threshold of FC and FDR.

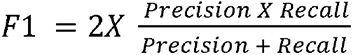

The G-mean score, or geometric mean, is a performance metric used in machine learning to evaluate the effectiveness of a classification model, particularly in the context of imbalanced datasets. It provides a balance between the sensitivity (true positive rate) and specificity (true negative rate) of the model. By taking the geometric mean of these two rates, the G-mean score ensures that a high value is only achieved when both sensitivity and specificity are high.

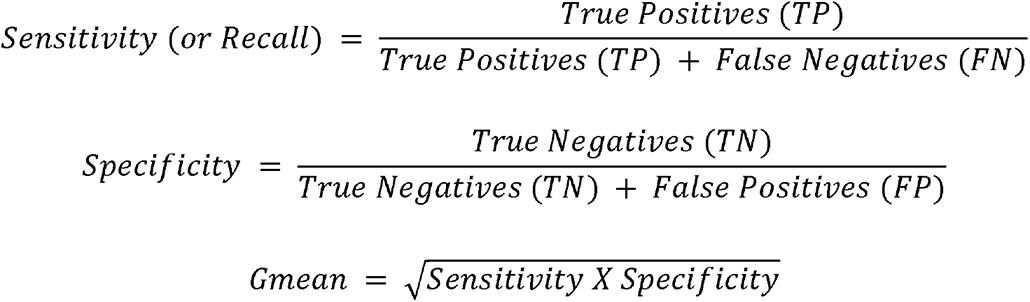

### MAGMa key functionalities

#### Row Normalization

For each PSM used in quantification, the row contains a relative intensity value for each TMT reporter tag. These values are normalized by dividing each by the average intensity of all reporter tags in that row. This step ensures that the ratios are maintained and consistent across the dataset.

#### Column Normalization

Column normalization is performed under the assumption that most proteins in our dataset are not significantly up- or down-regulated. We achieve this by using median ratio-based normalization, applying it to all peptides in the benchmarking dataset but only utilizing the peptides associated with human proteins for adjustment. Since human proteins are expected to exhibit a 1:1 ratio and remain unchanged across different reporter tags, they serve as a reliable reference for normalizing the yeast proteins.

#### Role of values separated across fractions

Herein, we test the assumption that the same peptides detected in different fractions (and thus in different machine runs, each subject to distinct run biases) can be treated as independent measurements for the same protein. To validate this, we examine the correlation between the raw measurements of identical peptides randomly sampled from two different fractions under the same experimental condition.

We base this assumption on the principle that values separated across fractions are independent. This is because these values were acquired in separate machine runs, and theoretically, a hydrophobicity gradient separation should result in each peptide appearing in only one fraction. Practically, this assumption is supported by our observation of a low correlation between raw intensity measurements for unique peptides found in multiple fractions (Supplementary Fig. 6 a,b).

#### MAGMa-PD implementation

If the proteomic study (using TMT or SILAC) prioritizes higher accuracy and precision over the number of identifications, we recommend using Proteome Discoverer’s quality metrics for assessing each PSM. Specifically, under the "Quan Info" column, remove all PSMs that do not qualify as high quality. This approach will improve the performance of MAGMa based on the metrics discussed in the manuscript (Fig. 2b,c,3b,c, Supplementary Fig. 2,3).

#### MAGMa-Limma implementation

This functionality is particularly crucial for processing SILAC datasets, where quantifications are naturally paired, requiring all subsequent analyses to account for this pairing. The key difference in this functionality lies in how we calculate the *P*-value and subsequently the FDR for each protein.

To achieve similar functionality for TMT datasets, we need to consider all possible ways of calculating ratios that are independent of each other, thus constituting independent trials. We perform this calculation per fraction per protein. For example, if a protein in the dataset (as illustrated in Supplementary Fig. 7a) had three replicates per condition and was found in two fractions, there would be 6x2 = 12 ways to obtain independent trials.

A simulation study was conducted to evaluate whether averaging *P*-values calculated for each independent trial of a protein is a viable approach for obtaining a representative *P*-value per protein. The feasibility of this method hinges on the assumption that the average of *P*-values will follow a uniform distribution, given that *P*-values under the null hypothesis should follow a uniform distribution, assuming all other conditions (e.g., the test statistic distribution) are met.

In this simulation, we considered a dataset with a fixed number of replicates (three in this case). The control values were drawn from a normal distribution with a mean of 0 and a standard deviation of 1, while the treatment values were drawn from a normal distribution with a mean of 0 and a variable standard deviation. Under the null hypothesis (no difference between treatment and control), we calculated the *P*-value for every possible independent ratio (six ways for three replicates per condition in one fraction, as explained in Supplementary Fig. 7a). These *P*-values were computed using the *Limma*^31^ package in R.

We then averaged these *P*-values and examined their distribution. The results showed that the averaged *P*-values approximate a uniform distribution (Supplementary Fig. 7b). However, this method is conservative and tends to yield more false negatives, as indicated by the scarcity of small *P*-values in the distribution.

#### PSM-ratio based methods

In this study, we retain all PSM information as quantitative representatives of the protein. Ratios for each PSM are calculated by pairing the replicates in one specific manner. For instance, if there are three replicates each for Wild-Type (WT) and Control (C), the pairings would be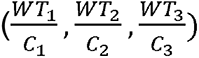. The fold change (FC) is determined as the median of all possible ratios associated with a protein. The Mann-Whitney U test is then performed, comparing each protein’s ratios to every other protein in the dataset to obtain raw p-values. These p-values are subsequently adjusted using the Benjamini-Hochberg correction to obtain the false discovery rate (FDR).

