## Supplementary Figures and Tables for "MAGMa: Your Comprehensive Tool for Differential Expression Analysis in Mass-Spectrometry Proteomic Data"

From Shagun Gupta *et al*.

### Supplementary Tables

(a)

| Tool | | All proteins | | | | Low abundant proteins | | | | | | | | High abundant proteins | | | |
|---|---|---|---|---|---|---|---|---|---|---|---|---|---|---|---|---|---|
| | | All PSMs | | | | <=2 PSMs | | | | <=3 PSMs | | | | >=5 PSMs | | | |
| | | G-mean | Recall | F1 | Accuracy | G-mean | Recall | F1 | Accuracy | G-mean | Recall | F1 | Accuracy | G-mean | Recall | F1 | Accuracy |
| COMET | Magma | **0.915** | **0.871** | **0.873** | **0.940** | **0.818** | **0.717** | **0.732** | **0.886** | **0.840** | **0.755** | **0.760** | **0.896** | **0.999** | **1.000** | **0.996** | **0.998** |
| | MSstatsTMT | 0.909 | 0.861 | 0.868 | 0.937 | 0.807 | 0.698 | 0.725 | 0.881 | 0.831 | 0.737 | 0.754 | 0.892 | 0.998 | **1.000** | 0.995 | 0.997 |
| | PSM-ratio | 0.761 | 0.583 | 0.727 | 0.895 | 0.392 | 0.156 | 0.261 | 0.802 | 0.490 | 0.243 | 0.381 | 0.822 | 0.993 | 0.987 | 0.991 | 0.995 |
| | Magma-Limma | 0.877 | 0.800 | 0.831 | 0.923 | 0.806 | 0.705 | 0.710 | 0.876 | 0.816 | 0.713 | 0.732 | 0.886 | 0.952 | 0.908 | 0.950 | 0.974 |
| SEQUEST | Magma | **0.931** | **0.897** | **0.898** | **0.950** | **0.846** | **0.768** | **0.770** | **0.894** | **0.864** | **0.794** | **0.796** | **0.908** | **0.994** | 0.990 | **0.991** | **0.995** |
| | MSstatsTMT | 0.928 | 0.890 | 0.895 | 0.948 | 0.840 | 0.757 | 0.767 | 0.890 | 0.858 | 0.781 | 0.792 | 0.904 | **0.994** | **0.992** | **0.991** | **0.995** |
| | PSM-ratio | 0.788 | 0.624 | 0.761 | 0.902 | 0.456 | 0.210 | 0.339 | 0.803 | 0.536 | 0.289 | 0.440 | 0.828 | 0.972 | 0.947 | 0.970 | 0.984 |
| | Magma-Limma | 0.901 | 0.837 | 0.868 | 0.937 | 0.856 | 0.784 | 0.783 | 0.900 | 0.860 | 0.782 | 0.795 | 0.909 | 0.947 | 0.901 | 0.943 | 0.969 |
| | Magma-PD | **0.973** | **0.963** | 0.957 | 0.979 | 0.930 | 0.894 | 0.889 | 0.952 | 0.939 | 0.907 | 0.904 | 0.959 | **0.996** | **0.997** | **0.992** | **0.996** |
| | PD 2.3 | **0.975** | **0.963** | **0.962** | **0.981** | **0.945** | **0.915** | **0.917** | **0.962** | **0.948** | **0.919** | **0.923** | **0.965** | 0.994 | 0.992 | 0.991 | 0.995 |
| MaxQuant | | 0.963 | 0.948 | 0.934 | 0.973 | 0.915 | 0.876 | 0.852 | 0.940 | 0.924 | 0.886 | 0.866 | 0.949 | 0.992 | 0.991 | 0.984 | 0.993 |
| MSFragger TMTIntegrator | | 0.927 | 0.879 | 0.916 | 0.941 | 0.883 | 0.805 | 0.858 | 0.917 | 0.892 | 0.820 | 0.871 | 0.921 | 0.963 | 0.939 | 0.961 | 0.967 |

(b)

| Tool | | All proteins | | | | Low abundant proteins | | | | | | | | High abundant proteins | | | |
|---|---|---|---|---|---|---|---|---|---|---|---|---|---|---|---|---|---|
| | | All PSMs | | | | <=2 PSMs | | | | <=3 PSMs | | | | >=5 PSMs | | | |
| | | G-mean | Recall | F1 | Accuracy | G-mean | Recall | F1 | Accuracy | G-mean | Recall | F1 | Accuracy | G-mean | Recall | F1 | Accuracy |
| COMET | Magma | **0.824** | **0.687** | **0.795** | **0.916** | **0.668** | **0.457** | **0.593** | **0.863** | **0.698** | **0.497** | **0.633** | **0.874** | **0.954** | **0.911** | **0.951** | **0.975** |
| | MSstatsTMT | 0.805 | 0.655 | 0.774 | 0.908 | 0.646 | 0.427 | 0.567 | 0.854 | 0.664 | 0.450 | 0.592 | 0.861 | 0.952 | 0.909 | 0.950 | 0.974 |
| | PSM-ratio | 0.725 | 0.528 | 0.682 | 0.882 | 0.371 | 0.139 | 0.236 | 0.798 | 0.448 | 0.203 | 0.329 | 0.813 | 0.960 | 0.924 | 0.958 | 0.978 |
| | Magma-Limma | 0.799 | 0.646 | 0.766 | 0.907 | 0.670 | 0.460 | 0.594 | 0.864 | 0.685 | 0.479 | 0.617 | 0.870 | 0.920 | 0.847 | 0.916 | 0.958 |
| SEQUEST | Magma | **0.842** | **0.717** | 0.818 | **0.921** | **0.701** | **0.505** | **0.634** | **0.865** | **0.717** | **0.526** | **0.657** | **0.875** | 0.949 | 0.902 | 0.946 | 0.971 |
| | MSstatsTMT | **0.842** | **0.717** | **0.820** | **0.921** | 0.686 | 0.482 | 0.619 | 0.858 | 0.706 | 0.509 | 0.647 | 0.870 | **0.959** | **0.922** | **0.957** | **0.977** |
| | PSM-ratio | 0.752 | 0.567 | 0.718 | 0.888 | 0.437 | 0.192 | 0.315 | 0.799 | 0.503 | 0.255 | 0.398 | 0.820 | 0.937 | 0.880 | 0.934 | 0.965 |
| | Magma-Limma | 0.818 | 0.677 | 0.792 | 0.913 | 0.692 | 0.491 | 0.625 | 0.865 | 0.700 | 0.500 | 0.638 | 0.873 | 0.923 | 0.853 | 0.920 | 0.958 |
| | Magma-PD | 0.929 | 0.867 | 0.920 | 0.964 | 0.845 | 0.722 | 0.817 | 0.930 | 0.843 | 0.718 | 0.818 | 0.931 | **0.987** | **0.975** | **0.986** | **0.992** |
| | PD 2.3 | **0.930** | **0.870** | **0.923** | 0.963 | **0.861** | **0.750** | **0.838** | 0.933 | 0.851 | 0.731 | 0.830 | 0.932 | 0.978 | 0.959 | 0.977 | 0.987 |
| MaxQuant | | 0.927 | 0.864 | 0.917 | **0.968** | 0.846 | 0.727 | 0.813 | **0.934** | **0.859** | **0.747** | **0.832** | **0.944** | 0.977 | 0.956 | 0.976 | 0.990 |
| MSFragger TMTIntegrator | | 0.814 | 0.669 | 0.794 | 0.874 | 0.701 | 0.497 | 0.655 | 0.838 | 0.725 | 0.530 | 0.686 | 0.842 | 0.921 | 0.858 | 0.916 | 0.932 |

(c )

| Tool | | All proteins | | | | Low abundant proteins | | | | | | | | High abundant proteins | | | |
|---|---|---|---|---|---|---|---|---|---|---|---|---|---|---|---|---|---|
| | | All PSMs | | | | <=2 PSMs | | | | <=3 PSMs | | | | >=5 PSMs | | | |
| | | G-mean | Recall | F1 | Accuracy | G-mean | Recall | F1 | Accuracy | G-mean | Recall | F1 | Accuracy | G-mean | Recall | F1 | Accuracy |
| COMET | Magma | **0.897** | **0.827** | 0.863 | **0.938** | **0.777** | **0.635** | 0.701 | **0.882** | **0.803** | **0.675** | 0.734 | **0.893** | 0.997 | **0.997** | 0.995 | 0.997 |
| | MSstatsTMT | 0.896 | 0.824 | **0.864** | **0.938** | 0.773 | 0.626 | **0.702** | 0.881 | 0.801 | 0.671 | **0.737** | **0.893** | **0.998** | **0.997** | **0.996** | **0.998** |
| | PSM-ratio | 0.697 | 0.486 | 0.652 | 0.875 | 0.189 | 0.036 | 0.069 | 0.782 | 0.305 | 0.093 | 0.170 | 0.795 | 0.985 | 0.972 | 0.983 | 0.991 |
| | Magma-Limma | 0.851 | 0.749 | 0.807 | 0.915 | 0.795 | 0.677 | 0.705 | 0.878 | 0.799 | 0.678 | 0.720 | 0.885 | 0.917 | 0.841 | 0.913 | 0.957 |
| SEQUEST | Magma | **0.918** | **0.860** | **0.893** | **0.949** | 0.812 | 0.690 | 0.750 | 0.893 | 0.830 | 0.716 | 0.775 | 0.906 | **0.994** | **0.990** | **0.991** | **0.995** |
| | MSstatsTMT | 0.915 | 0.858 | 0.888 | 0.946 | 0.817 | 0.702 | 0.755 | 0.891 | 0.832 | 0.722 | 0.776 | 0.903 | 0.992 | 0.987 | 0.989 | 0.994 |
| | PSM-ratio | 0.708 | 0.502 | 0.666 | 0.874 | 0.209 | 0.044 | 0.083 | 0.768 | 0.317 | 0.100 | 0.181 | 0.787 | 0.941 | 0.887 | 0.939 | 0.968 |
| | Magma-Limma | 0.890 | 0.817 | 0.856 | 0.933 | **0.851** | **0.775** | **0.777** | **0.898** | **0.853** | **0.769** | **0.787** | **0.907** | 0.931 | 0.871 | 0.926 | 0.961 |
| | Magma-PD | **0.967** | **0.947** | **0.954** | **0.978** | 0.908 | 0.848 | 0.870 | 0.946 | 0.920 | **0.865** | 0.891 | 0.955 | **0.997** | **0.997** | **0.994** | **0.996** |
| | PD 2.3 | 0.962 | 0.934 | 0.953 | 0.977 | **0.919** | **0.858** | **0.899** | **0.955** | **0.923** | 0.863 | **0.906** | **0.959** | 0.989 | 0.982 | 0.986 | 0.992 |
| MaxQuant | | 0.959 | 0.930 | 0.943 | 0.977 | 0.899 | 0.830 | 0.859 | 0.946 | 0.912 | 0.849 | 0.876 | 0.955 | 0.992 | 0.988 | 0.988 | 0.995 |
| MSFragger TMTIntegrator | | 0.917 | 0.855 | 0.908 | 0.937 | 0.871 | 0.777 | 0.850 | 0.915 | 0.879 | 0.790 | 0.861 | 0.918 | 0.961 | 0.935 | 0.959 | 0.965 |

Supplementary Table 1: **Performance evaluation of open-source and proprietary software(s) with MAGMa on the in-house benchmarking dataset on comparison with expected FC of 10 for yeast proteins and 1 for human proteins.** (a) Threshold of FC:2, FDR: 5% (b) Threshold of FC:5, FDR: 5% (c) Threshold of FC:2, FDR: 1%.

(a)

| Tool | | All proteins | | | | Low abundant proteins | | | | | | | | High abundant proteins | | | |
|---|---|---|---|---|---|---|---|---|---|---|---|---|---|---|---|---|---|
| | | **All PSMs** | | | | **<=2 PSMs** | | | | **<=3 PSMs** | | | | **>=5 PSMs** | | | |
| | | G-mean | Recall | F1 | Accuracy | G-mean | Recall | F1 | Accuracy | G-mean | Recall | F1 | Accuracy | G-mean | Recall | F1 | Accuracy |
| COMET | Magma | **0.821** | **0.684** | **0.793** | **0.915** | **0.640** | **0.419** | **0.557** | **0.854** | **0.669** | **0.458** | **0.596** | **0.864** | **0.976** | **0.954** | **0.975** | **0.987** |
| | MSstatsTMT | 0.800 | 0.647 | 0.770 | 0.907 | 0.617 | 0.388 | 0.533 | 0.847 | 0.647 | 0.426 | 0.573 | 0.858 | 0.962 | 0.926 | 0.961 | 0.980 |
| | PSM-ratio | 0.736 | 0.544 | 0.698 | 0.887 | 0.382 | 0.147 | 0.251 | 0.804 | 0.459 | 0.212 | 0.343 | 0.818 | 0.975 | 0.952 | 0.974 | 0.986 |
| | Magma-Limma | 0.489 | 0.241 | 0.381 | 0.814 | 0.458 | 0.213 | 0.335 | 0.818 | 0.450 | 0.205 | 0.327 | 0.815 | 0.540 | 0.292 | 0.451 | 0.808 |
| SEQUEST | Magma | **0.862** | **0.751** | **0.843** | **0.931** | **0.719** | **0.529** | **0.659** | **0.873** | **0.730** | **0.544** | **0.676** | **0.882** | **0.977** | **0.955** | **0.976** | **0.987** |
| | MSstatsTMT | 0.851 | 0.732 | 0.830 | 0.925 | 0.699 | 0.500 | 0.636 | 0.863 | 0.720 | 0.528 | 0.665 | 0.876 | 0.966 | 0.935 | 0.964 | 0.980 |
| | PSM-ratio | 0.738 | 0.547 | 0.703 | 0.885 | 0.358 | 0.129 | 0.224 | 0.787 | 0.426 | 0.182 | 0.304 | 0.805 | 0.951 | 0.905 | 0.949 | 0.973 |
| | Magma-Limma | 0.685 | 0.473 | 0.632 | 0.865 | 0.615 | 0.386 | 0.533 | 0.845 | 0.611 | 0.379 | 0.530 | 0.849 | 0.758 | 0.575 | 0.730 | 0.880 |
| | Magma-PD | **0.934** | **0.876** | **0.927** | **0.966** | **0.845** | **0.722** | **0.819** | **0.931** | **0.846** | **0.722** | **0.824** | **0.934** | **0.989** | **0.980** | **0.988** | **0.994** |
| | PD 2.3 | 0.918 | 0.846 | 0.910 | 0.958 | 0.817 | 0.676 | 0.788 | 0.915 | 0.823 | 0.684 | 0.798 | 0.921 | 0.972 | 0.947 | 0.971 | 0.984 |
| MaxQuant | | 0.918 | 0.846 | 0.910 | **0.966** | 0.829 | 0.694 | 0.798 | **0.931** | 0.841 | 0.713 | 0.815 | 0.940 | 0.975 | 0.950 | 0.974 | 0.989 |
| MSFragger TMTIntegrator | | 0.885 | 0.793 | 0.874 | 0.916 | 0.810 | 0.668 | 0.783 | 0.886 | 0.825 | 0.691 | 0.803 | 0.890 | 0.953 | 0.920 | 0.950 | 0.959 |

(b)

| Tool | | All proteins | | | | Low abundant proteins | | | | | | | | High abundant proteins | | | |
|---|---|---|---|---|---|---|---|---|---|---|---|---|---|---|---|---|---|
| | | **All PSMs** | | | | **<=2 PSMs** | | | | **<=3 PSMs** | | | | **>=5 PSMs** | | | |
| | | G-mean | Recall | F1 | Accuracy | G-mean | Recall | F1 | Accuracy | G-mean | Recall | F1 | Accuracy | G-mean | Recall | F1 | Accuracy |
| COMET | Magma | **0.720** | **0.523** | **0.675** | **0.880** | **0.536** | **0.292** | **0.432** | **0.833** | **0.559** | **0.317** | **0.463** | **0.839** | 0.879 | 0.774 | 0.871 | 0.939 |
| | MSstatsTMT | 0.654 | 0.431 | 0.592 | 0.857 | 0.503 | 0.257 | 0.391 | 0.821 | 0.511 | 0.265 | 0.403 | 0.824 | 0.807 | 0.652 | 0.788 | 0.906 |
| | PSM-ratio | 0.681 | 0.466 | 0.630 | 0.869 | 0.360 | 0.130 | 0.225 | 0.800 | 0.426 | 0.182 | 0.302 | 0.812 | **0.905** | **0.820** | **0.900** | **0.951** |
| | Magma-Limma | 0.482 | 0.234 | 0.372 | 0.812 | 0.450 | 0.205 | 0.326 | 0.817 | 0.438 | 0.195 | 0.314 | 0.814 | 0.537 | 0.289 | 0.448 | 0.807 |
| SEQUEST | Magma | **0.773** | **0.602** | **0.741** | **0.897** | **0.602** | **0.368** | **0.517** | **0.841** | **0.607** | **0.374** | **0.526** | **0.847** | **0.905** | **0.819** | **0.901** | **0.949** |
| | MSstatsTMT | 0.761 | 0.585 | 0.726 | 0.890 | 0.577 | 0.339 | 0.485 | 0.828 | 0.588 | 0.351 | 0.502 | 0.837 | 0.901 | 0.812 | 0.895 | 0.947 |
| | PSM-ratio | 0.694 | 0.482 | 0.647 | 0.869 | 0.346 | 0.120 | 0.211 | 0.785 | 0.402 | 0.163 | 0.276 | 0.801 | 0.894 | 0.799 | 0.888 | 0.944 |
| | Magma-Limma | 0.652 | 0.428 | 0.590 | 0.854 | 0.556 | 0.315 | 0.458 | 0.829 | 0.551 | 0.308 | 0.454 | 0.833 | 0.741 | 0.549 | 0.709 | 0.873 |
| | Magma-PD | **0.876** | **0.769** | **0.866** | **0.942** | **0.759** | **0.581** | **0.723** | **0.904** | **0.756** | **0.575** | **0.722** | **0.905** | **0.950** | **0.903** | **0.949** | **0.973** |
| | PD 2.3 | 0.862 | 0.746 | 0.851 | 0.934 | 0.751 | 0.568 | 0.712 | 0.893 | 0.743 | 0.556 | 0.705 | 0.894 | 0.929 | 0.863 | 0.926 | 0.961 |
| MaxQuant | | 0.824 | 0.680 | 0.805 | 0.933 | 0.713 | 0.513 | 0.664 | 0.899 | 0.736 | 0.545 | 0.695 | 0.911 | 0.895 | 0.802 | 0.890 | 0.957 |
| MSFragger TMTIntegrator | | 0.768 | 0.594 | 0.739 | 0.847 | 0.639 | 0.411 | 0.576 | 0.812 | 0.663 | 0.442 | 0.607 | 0.815 | 0.892 | 0.805 | 0.884 | 0.909 |

(c )

| Tool | | All proteins | | | | Low abundant proteins | | | | | | | | High abundant proteins | | | |
|---|---|---|---|---|---|---|---|---|---|---|---|---|---|---|---|---|---|
| | | **All PSMs** | | | | **<=2 PSMs** | | | | **<=3 PSMs** | | | | **>=5 PSMs** | | | |
| | | G-mean | Recall | F1 | Accuracy | G-mean | Recall | F1 | Accuracy | G-mean | Recall | F1 | Accuracy | G-mean | Recall | F1 | Accuracy |
| COMET | Magma | **0.782** | **0.618** | **0.750** | **0.902** | **0.547** | 0.304 | 0.445 | **0.834** | **0.581** | 0.342 | 0.489 | **0.843** | **0.974** | **0.949** | **0.973** | **0.986** |
| | MSstatsTMT | 0.771 | 0.600 | 0.737 | 0.898 | **0.547** | **0.305** | **0.448** | 0.832 | **0.581** | **0.343** | **0.492** | 0.842 | 0.962 | 0.926 | 0.961 | 0.980 |
| | PSM-ratio | 0.681 | 0.464 | 0.632 | 0.871 | 0.208 | 0.043 | 0.083 | 0.785 | 0.307 | 0.094 | 0.171 | 0.796 | 0.965 | 0.931 | 0.963 | 0.981 |
| | Magma-Limma | 0.000 | 0.000 | 0.000 | 0.000 | 0.000 | 0.000 | 0.000 | 0.000 | 0.000 | 0.000 | 0.000 | 0.000 | 0.000 | 0.000 | 0.000 | 0.000 |
| SEQUEST | Magma | **0.814** | **0.668** | **0.789** | **0.912** | **0.624** | **0.396** | **0.545** | **0.846** | **0.640** | **0.415** | **0.567** | **0.856** | **0.953** | **0.910** | **0.951** | **0.974** |
| | MSstatsTMT | 0.805 | 0.654 | 0.778 | 0.907 | 0.616 | 0.386 | 0.534 | 0.839 | 0.636 | 0.410 | 0.562 | 0.851 | 0.944 | 0.892 | 0.941 | 0.968 |
| | PSM-ratio | 0.675 | 0.456 | 0.626 | 0.864 | 0.188 | 0.035 | 0.068 | 0.769 | 0.281 | 0.079 | 0.146 | 0.785 | 0.912 | 0.832 | 0.908 | 0.953 |
| | Magma-Limma | 0.000 | 0.000 | 0.000 | 0.000 | 0.000 | 0.000 | 0.000 | 0.000 | 0.000 | 0.000 | 0.000 | 0.000 | 0.000 | 0.000 | 0.000 | 0.000 |
| | Magma-PD | **0.892** | **0.798** | **0.882** | 0.948 | **0.755** | **0.576** | **0.715** | 0.901 | **0.761** | 0.583 | 0.724 | 0.905 | **0.969** | **0.939** | **0.967** | **0.982** |
| | PD 2.3 | 0.847 | 0.720 | 0.834 | 0.928 | 0.696 | 0.489 | 0.644 | 0.875 | 0.699 | 0.491 | 0.650 | 0.879 | 0.926 | 0.858 | 0.923 | 0.960 |
| MaxQuant | | 0.886 | 0.788 | 0.876 | **0.955** | 0.741 | 0.554 | 0.697 | **0.906** | 0.771 | **0.598** | **0.736** | **0.920** | 0.965 | 0.932 | 0.965 | 0.985 |
| MSFragger TMTIntegrator | | 0.828 | 0.692 | 0.810 | 0.881 | 0.715 | 0.517 | 0.670 | 0.842 | 0.732 | 0.541 | 0.693 | 0.844 | 0.937 | 0.889 | 0.934 | 0.945 |

Supplementary Table 2: **Performance evaluation of open-source and proprietary software(s) with MAGMa on the in-house benchmarking dataset on comparison with expected FC of 4 for yeast proteins and 1 for human proteins.** (a) Threshold of FC:2, FDR: 5% (b) Threshold of FC:3, FDR: 5% (c) Threshold of FC:2, FDR: 1%.

| | | All proteins | | | | Low abundant proteins | | | | | | | | High abundant proteins | | | |
|---|---|---|---|---|---|---|---|---|---|---|---|---|---|---|---|---|---|
| **Tool** | | **All PSMs** | | | | **<=2 PSMs** | | | | **<=3 PSMs** | | | | **>=5 PSMs** | | | |
| | | G-mean | Recall | F1 | Accuracy | G-mean | Recall | F1 | Accuracy | G-mean | Recall | F1 | Accuracy | G-mean | Recall | F1 | Accuracy |
| COMET | Magma | 0.919 | 0.877 | 0.879 | **0.944** | 0.857 | 0.793 | 0.794 | 0.891 | 0.889 | 0.836 | 0.840 | 0.918 | 0.990 | 0.980 | 0.990 | 0.997 |
| | MSstatsTMT | **0.922** | **0.883** | 0.879 | **0.944** | 0.863 | 0.807 | 0.797 | 0.893 | 0.893 | 0.844 | 0.843 | 0.919 | **0.999** | **1.000** | **0.995** | **0.998** |
| | PSM-ratio | 0.840 | 0.714 | 0.814 | 0.924 | 0.676 | 0.466 | 0.615 | 0.845 | 0.772 | 0.605 | 0.735 | 0.888 | 0.994 | 0.990 | 0.990 | 0.997 |
| | Magma-Limma | 0.920 | 0.879 | **0.880** | **0.944** | **0.875** | **0.829** | **0.811** | **0.898** | **0.899** | **0.856** | **0.850** | **0.922** | 0.975 | 0.951 | 0.975 | 0.992 |
| SEQUEST | Magma | 0.954 | 0.931 | 0.930 | 0.968 | 0.916 | 0.878 | 0.878 | 0.935 | 0.938 | 0.909 | 0.908 | 0.952 | **1.000** | **1.000** | **1.000** | **1.000** |
| | MSstatsTMT | 0.952 | 0.926 | 0.928 | 0.968 | 0.913 | 0.869 | 0.876 | 0.936 | 0.937 | 0.905 | 0.908 | **0.954** | 0.992 | 0.986 | 0.986 | 0.996 |
| | PSM-ratio | 0.859 | 0.743 | 0.841 | 0.937 | 0.710 | 0.510 | 0.662 | 0.864 | 0.807 | 0.658 | 0.779 | 0.906 | **1.000** | **1.000** | **1.000** | **1.000** |
| | Magma-Limma | **0.956** | **0.934** | **0.932** | **0.969** | **0.922** | **0.888** | **0.886** | **0.938** | **0.940** | **0.912** | **0.911** | **0.954** | **1.000** | **1.000** | **1.000** | **1.000** |
| | Magma-PD | **0.978** | **0.970** | **0.963** | **0.983** | **0.975** | **0.978** | **0.957** | **0.975** | **0.980** | **0.975** | **0.969** | **0.983** | 0.984 | 0.977 | 0.966 | 0.989 |
| | PD 2.3 | 0.967 | 0.943 | 0.955 | 0.980 | 0.963 | 0.947 | 0.947 | 0.970 | 0.973 | 0.958 | 0.965 | 0.981 | 0.972 | 0.953 | 0.953 | 0.984 |
| MaxQuant | | 0.940 | 0.907 | 0.908 | 0.959 | 0.905 | 0.863 | 0.863 | 0.926 | 0.918 | 0.879 | 0.879 | 0.938 | **0.994** | **0.988** | **0.994** | **0.998** |
| MSFragger TMTIntegrator | | 0.911 | 0.855 | 0.875 | 0.945 | 0.874 | 0.812 | 0.824 | 0.906 | 0.901 | 0.850 | 0.857 | 0.928 | 0.930 | 0.864 | 0.927 | 0.978 |

Supplementary Table 3: Comparison analysis on published phospho-proteome benchmarking dataset on 10-1 comparison at FC:2, FDR: 5%

| | | All proteins | | | | Low abundant proteins | | | | | | | | High abundant proteins | | | |
|---|---|---|---|---|---|---|---|---|---|---|---|---|---|---|---|---|---|
| **Tool** | | **All PSMs** | | | | **<=2 PSMs** | | | | **<=3 PSMs** | | | | **>=5 PSMs** | | | |
| | | G-mean | Recall | F1 | Accuracy | G-mean | Recall | F1 | Accuracy | G-mean | Recall | F1 | Accuracy | G-mean | Recall | F1 | Accuracy |
| COMET | Magma | 0.802 | 0.648 | 0.775 | 0.913 | 0.672 | 0.459 | 0.611 | 0.845 | 0.747 | 0.564 | 0.707 | 0.879 | 0.929 | 0.863 | 0.926 | 0.977 |
| | MSstatsTMT | **0.818** | **0.675** | **0.793** | **0.918** | **0.706** | **0.506** | **0.653** | **0.860** | **0.767** | **0.596** | **0.730** | **0.886** | 0.938 | 0.882 | 0.933 | 0.978 |
| | PSM-ratio | 0.761 | 0.581 | 0.730 | 0.901 | 0.553 | 0.308 | 0.465 | 0.812 | 0.675 | 0.458 | 0.623 | 0.857 | **0.945** | **0.892** | **0.943** | **0.982** |
| | Magma-Limma | 0.798 | 0.642 | 0.767 | 0.910 | 0.699 | 0.500 | 0.641 | 0.851 | 0.747 | 0.567 | 0.704 | 0.877 | 0.924 | 0.853 | 0.921 | 0.975 |
| SEQUEST | Magma | **0.855** | **0.734** | **0.839** | **0.935** | 0.767 | 0.595 | 0.733 | 0.884 | 0.820 | 0.677 | 0.798 | 0.911 | **0.959** | **0.919** | **0.958** | **0.987** |
| | MSstatsTMT | 0.847 | 0.721 | 0.831 | 0.934 | **0.786** | **0.623** | **0.756** | **0.896** | **0.823** | **0.682** | **0.801** | **0.915** | 0.923 | 0.851 | 0.920 | 0.975 |
| | PSM-ratio | 0.794 | 0.632 | 0.771 | 0.915 | 0.632 | 0.401 | 0.566 | 0.840 | 0.737 | 0.546 | 0.701 | 0.883 | 0.952 | 0.905 | 0.950 | 0.984 |
| | Magma-Limma | 0.847 | 0.721 | 0.831 | 0.932 | 0.761 | 0.585 | 0.725 | 0.881 | 0.814 | 0.667 | 0.790 | 0.908 | 0.952 | 0.905 | 0.950 | 0.984 |
| | Magma-PD | **0.870** | **0.759** | **0.860** | **0.945** | **0.816** | **0.667** | **0.800** | **0.904** | **0.880** | **0.775** | **0.873** | **0.938** | 0.877 | 0.773 | 0.861 | 0.958 |
| | PD 2.3 | 0.850 | 0.724 | 0.836 | 0.937 | 0.761 | 0.579 | 0.733 | 0.881 | 0.848 | 0.718 | 0.836 | 0.923 | 0.874 | 0.767 | 0.857 | 0.957 |
| MaxQuant | | 0.866 | 0.758 | 0.844 | 0.937 | 0.802 | 0.657 | 0.765 | 0.891 | 0.833 | 0.706 | 0.803 | 0.911 | **0.951** | **0.904** | **0.949** | **0.986** |
| MSFragger TMTIntegrator | | 0.789 | 0.627 | 0.758 | 0.911 | 0.677 | 0.465 | 0.618 | 0.844 | 0.753 | 0.573 | 0.713 | 0.882 | 0.875 | 0.765 | 0.867 | 0.962 |

Supplementary Table 4: Comparison analysis on published phospho-proteome benchmarking dataset on 4-1 comparison at FC:2, FDR: 5%

| Tool | | All proteins | | | | Low abundant proteins | | | | | | | | High abundant proteins | | | |
|---|---|---|---|---|---|---|---|---|---|---|---|---|---|---|---|---|---|
| | | **All PSMs** | | | | **<=2 PSMs** | | | | **<=3 PSMs** | | | | **>=5 PSMs** | | | |
| | | G-mean | Recall | F1 | Accuracy | G-mean | Recall | F1 | Accuracy | G-mean | Recall | F1 | Accuracy | G-mean | Recall | F1 | Accuracy |
| COMET | Magma | **0.915** | **0.871** | **0.873** | **0.940** | **0.818** | **0.717** | **0.732** | **0.886** | **0.840** | **0.755** | **0.760** | **0.896** | **0.999** | **1.000** | 0.996 | 0.998 |
| | - ROW NORM | 0.862 | 0.767 | 0.823 | 0.922 | 0.779 | 0.645 | 0.695 | 0.877 | 0.783 | 0.646 | 0.706 | 0.883 | 0.971 | 0.944 | 0.969 | 0.984 |
| | - COLUMN NORM | 0.911 | 0.861 | 0.871 | **0.940** | 0.807 | 0.696 | 0.720 | 0.882 | 0.832 | 0.738 | 0.755 | 0.895 | **0.999** | **1.000** | **0.997** | **0.999** |
| | - FRACTION | 0.904 | 0.847 | 0.864 | 0.937 | 0.799 | 0.684 | 0.711 | 0.879 | 0.820 | 0.715 | 0.740 | 0.890 | 0.998 | 0.997 | 0.996 | 0.998 |
| | - ROW NORM - FRACTION | 0.882 | 0.812 | 0.834 | 0.923 | 0.790 | 0.668 | 0.701 | 0.876 | 0.798 | 0.679 | 0.714 | 0.881 | 0.977 | 0.967 | 0.966 | 0.982 |
| SEQUEST | Magma | **0.931** | **0.897** | 0.898 | 0.950 | **0.846** | **0.768** | **0.770** | 0.894 | **0.864** | **0.794** | 0.796 | 0.908 | **0.994** | 0.990 | **0.991** | **0.995** |
| | - ROW NORM | 0.872 | 0.779 | 0.841 | 0.928 | 0.804 | 0.681 | 0.736 | 0.887 | 0.800 | 0.668 | 0.737 | 0.892 | 0.941 | 0.887 | 0.938 | 0.967 |
| | - COLUMN NORM | 0.928 | 0.884 | **0.901** | **0.952** | 0.836 | 0.740 | **0.770** | **0.898** | 0.858 | 0.773 | **0.800** | **0.912** | **0.994** | 0.990 | **0.991** | **0.995** |
| | - FRACTION | 0.926 | 0.887 | 0.893 | 0.948 | 0.838 | 0.752 | 0.762 | 0.891 | 0.854 | 0.773 | 0.786 | 0.905 | **0.994** | **0.992** | **0.991** | **0.995** |
| | - ROW NORM - FRACTION | 0.908 | 0.858 | 0.867 | 0.936 | 0.829 | 0.737 | 0.752 | 0.888 | 0.839 | 0.750 | 0.767 | 0.896 | 0.975 | 0.965 | 0.964 | 0.980 |

Supplementary Table 5: Ablation study on MAGMa on in-house benchmarking dataset on 10-1 comparison at FC:2, FDR: 5%.

| Tool | | All proteins | | | | Low abundant proteins | | | | | | | | High abundant proteins | | | |
|---|---|---|---|---|---|---|---|---|---|---|---|---|---|---|---|---|---|
| | | **All PSMs** | | | | **<=2 PSMs** | | | | **<=3 PSMs** | | | | **>=5 PSMs** | | | |
| | | G-mean | Recall | F1 | Accuracy | G-mean | Recall | F1 | Accuracy | G-mean | Recall | F1 | Accuracy | G-mean | Recall | F1 | Accuracy |
| COMET | Magma | **0.821** | **0.684** | **0.793** | **0.915** | **0.640** | **0.419** | **0.557** | **0.854** | **0.669** | 0.458 | 0.596 | 0.864 | **0.976** | **0.954** | **0.975** | **0.987** |
| | No Row Norm | 0.659 | 0.439 | 0.595 | 0.858 | 0.566 | 0.327 | 0.465 | 0.835 | 0.556 | 0.315 | 0.455 | 0.835 | 0.788 | 0.622 | 0.766 | 0.898 |
| | No Column Norm | 0.782 | 0.618 | 0.750 | 0.902 | 0.563 | 0.322 | 0.465 | 0.838 | 0.598 | 0.363 | 0.512 | 0.848 | 0.962 | 0.926 | 0.961 | 0.980 |
| | No Fraction Separation | 0.720 | 0.523 | 0.675 | 0.881 | 0.514 | 0.269 | 0.406 | 0.829 | 0.535 | 0.290 | 0.433 | 0.834 | 0.904 | 0.817 | 0.898 | 0.950 |
| | No Row + No Fraction | 0.597 | 0.360 | 0.520 | 0.842 | 0.482 | 0.235 | 0.366 | 0.822 | 0.482 | **0.525** | **0.688** | **0.872** | 0.724 | 0.525 | 0.688 | 0.872 |
| SEQUEST | Magma | **0.862** | **0.751** | **0.843** | **0.931** | 0.719 | 0.529 | 0.659 | **0.873** | 0.730 | 0.544 | 0.676 | **0.882** | **0.977** | **0.955** | **0.976** | **0.987** |
| | No Row Norm | 0.713 | 0.513 | 0.665 | 0.873 | 0.637 | 0.415 | 0.557 | 0.847 | 0.608 | 0.376 | 0.523 | 0.844 | 0.811 | 0.658 | 0.794 | 0.904 |
| | No Column Norm | 0.821 | 0.679 | 0.797 | 0.915 | 0.634 | 0.409 | 0.557 | 0.849 | 0.653 | 0.433 | 0.584 | 0.860 | 0.955 | 0.912 | 0.953 | 0.975 |
| | No Fraction Separation | 0.859 | 0.746 | 0.839 | 0.930 | **0.721** | **0.533** | **0.660** | **0.873** | **0.732** | **0.546** | **0.677** | **0.882** | 0.969 | 0.940 | 0.968 | 0.982 |
| | No Row + No Fraction | 0.800 | 0.649 | 0.768 | 0.904 | 0.700 | 0.502 | 0.635 | 0.867 | 0.697 | 0.497 | 0.633 | 0.869 | 0.888 | 0.791 | 0.880 | 0.939 |

Supplementary Table 6: Ablation study on MAGMa on in-house benchmarking dataset on 4-1 comparison at FC:2, FDR: 5%.

| | | All proteins | | | | Low abundant proteins | | | | | | | | High abundant proteins | | | |
|---|---|---|---|---|---|---|---|---|---|---|---|---|---|---|---|---|---|
| **Tool** | | **All PSMs** | | | | **<=2 PSMs** | | | | **<=3 PSMs** | | | | **>=5 PSMs** | | | |
| | | G-mean | Recall | F1 | Accuracy | G-mean | Recall | F1 | Accuracy | G-mean | Recall | F1 | Accuracy | G-mean | Recall | F1 | Accuracy |
| COMET | MAGMa | **0.992** | **0.915** | 0.858 | 0.925 | **0.846** | **0.824** | **0.732** | 0.857 | **0.864** | **0.847** | **0.759** | 0.873 | 0.998 | **1.000** | 0.994 | 0.997 |
| | MSstatsTMT | 0.907 | 0.860 | **0.863** | **0.932** | 0.817 | 0.725 | **0.732** | **0.873** | 0.834 | 0.752 | 0.755 | **0.885** | **0.999** | **1.000** | **0.997** | **0.999** |
| | PSM-ratio | 0.770 | 0.597 | 0.739 | 0.896 | 0.453 | 0.207 | 0.335 | 0.803 | 0.535 | 0.289 | 0.439 | 0.825 | 0.993 | 0.987 | 0.991 | 0.995 |
| | MAGMa-Limma | 0.771 | 0.607 | 0.726 | 0.886 | 0.716 | 0.535 | 0.640 | 0.857 | 0.723 | 0.543 | 0.654 | 0.864 | 0.847 | 0.719 | 0.835 | 0.923 |
| SEQUEST | MAGMa | **0.931** | **0.910** | **0.889** | 0.941 | **0.851** | **0.805** | 0.766 | 0.875 | **0.869** | **0.825** | **0.791** | 0.893 | **0.996** | **0.997** | **0.991** | **0.995** |
| | MSstatsTMT | 0.923 | 0.883 | **0.889** | **0.943** | 0.837 | 0.757 | **0.768** | **0.883** | 0.854 | 0.779 | 0.790 | **0.898** | 0.993 | 0.993 | 0.989 | 0.994 |
| | PSM-ratio | 0.780 | 0.610 | 0.754 | 0.897 | 0.452 | 0.206 | 0.337 | 0.792 | 0.534 | 0.286 | 0.440 | 0.819 | 0.972 | 0.945 | 0.970 | 0.984 |
| | MAGMa-Limma | 0.813 | 0.673 | 0.779 | 0.902 | 0.757 | 0.597 | 0.695 | 0.866 | 0.764 | 0.605 | 0.708 | 0.877 | 0.864 | 0.748 | 0.853 | 0.928 |

Supplementary Table 7: Comparison analysis on in-house whole proteome benchmarking dataset on the ability to detect very small shifts in protein quantifications using empirical bayes moderation.

### Supplementary Figures

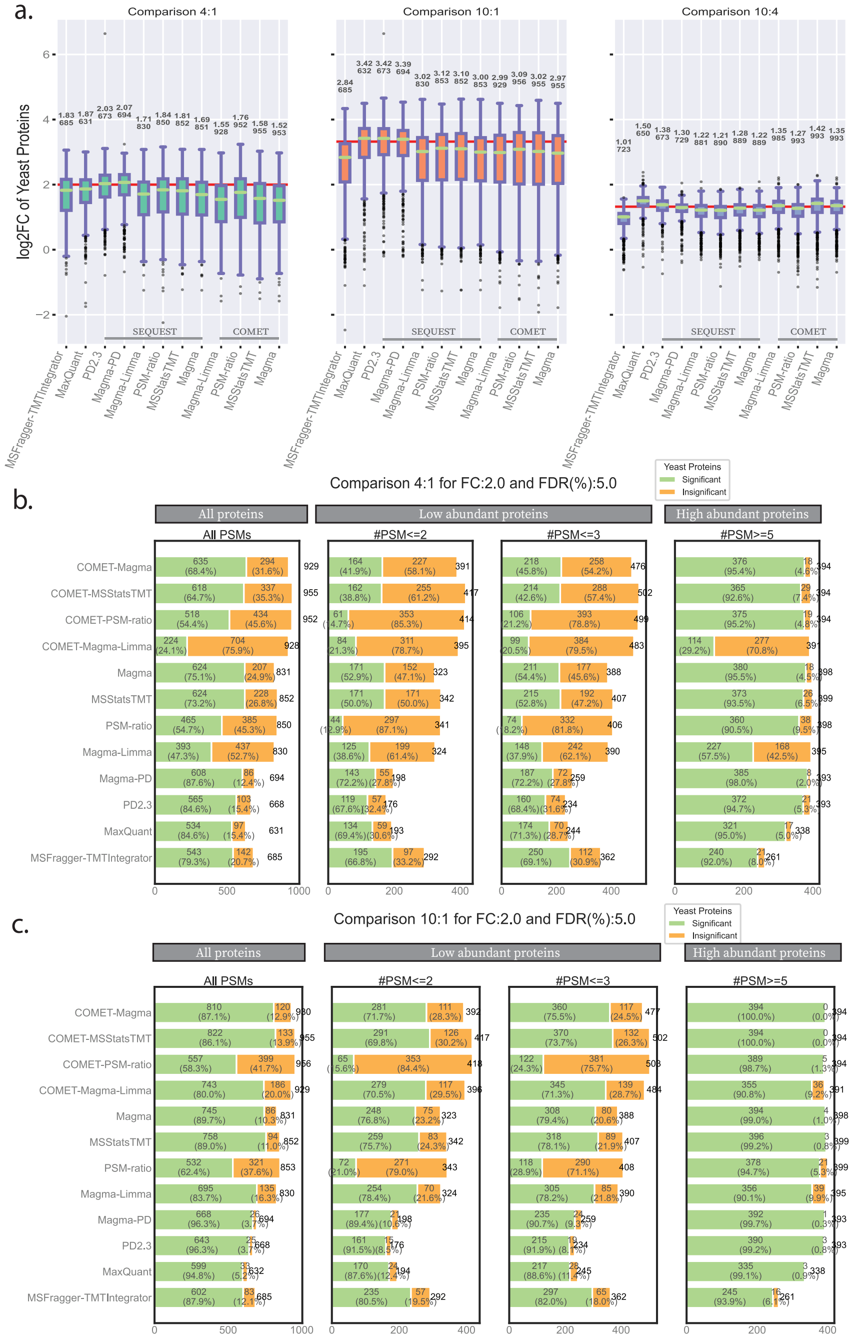


Supplementary Figure 1: **Performance evaluation of open-source and proprietary software(s) with MAGMa on the in-house benchmarking dataset.** (a) Distribution of log-2 transformed FC values for yeast proteins quantified with different combination of search and subsequent differential expression analysis tools. The three panels represent the three comparisons made (4:1, 10:1 and 10:4). The red line in each panel indicates the expected FC for that comparison. (b) Performance evaluation of various tools specifically on yeast proteins. It focuses on the tools' ability to detect an expected fold change (FC) of 4, comparing results across both low and highly abundant proteins. For each bar, in green are the number and proportion of yeast proteins that pass FC: 2 and FDR: 5% thresholds and in orange are number and proportion of yeast proteins that do not pass these thresholds. The number at the top of each bar is the total number of yeast proteins (c) Similar performance evaluation as (b) on yeast proteins on a comparison with an expected fold change (FC) of 10, comparing results across both low and highly abundant proteins.

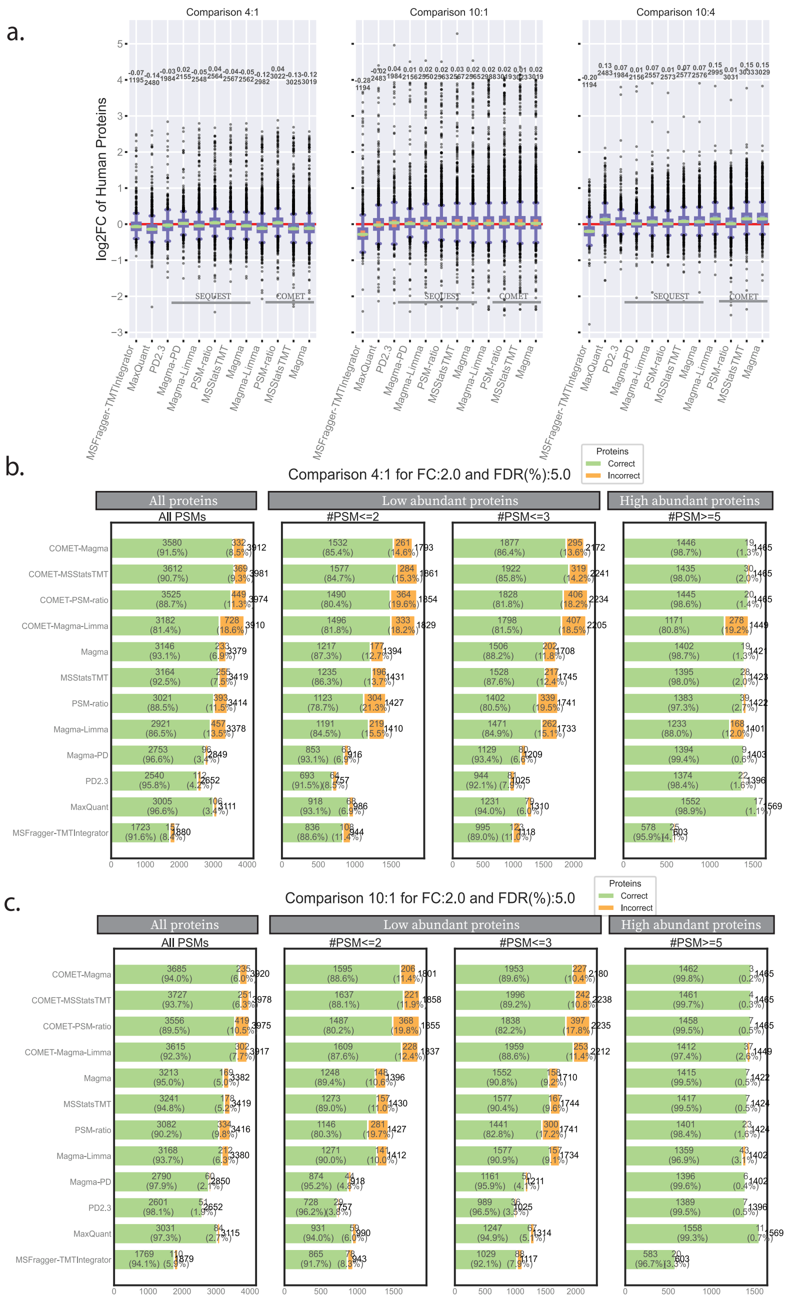


Supplementary Figure 2: **Performance evaluation of open-source and proprietary software(s) with MAGMa on the in-house benchmarking dataset.** (a) Distribution of log-2 transformed FC values for all human quantified with different combination of search and subsequent differential expression analysis tools. The three panels represent the three comparisons made (4:1, 10:1 and 10:4). The red line in each panel indicates the expected FC for that comparison. (b) Performance evaluation of tools on all proteins (yeast and human) on their ability to detect an expected fold change (FC) of 4 (yeast) and 1 (human), comparing results across both low and highly abundant proteins. For each bar, in green are the number and proportion of yeast proteins that pass FC: 2 and FDR: 5% thresholds and human proteins that do not pass these thresholds (correct calls). In orange are number and proportion of yeast proteins that do not pass these thresholds and number of human proteins that pass these thresholds (incorrect calls). The number at the top of each bar is the total number of proteins. (c) Similar performance evaluation as (b) on yeast and human proteins on a comparison with an expected fold change (FC) of 10 (yeast) and 1(human), comparing results across both low and highly abundant proteins.

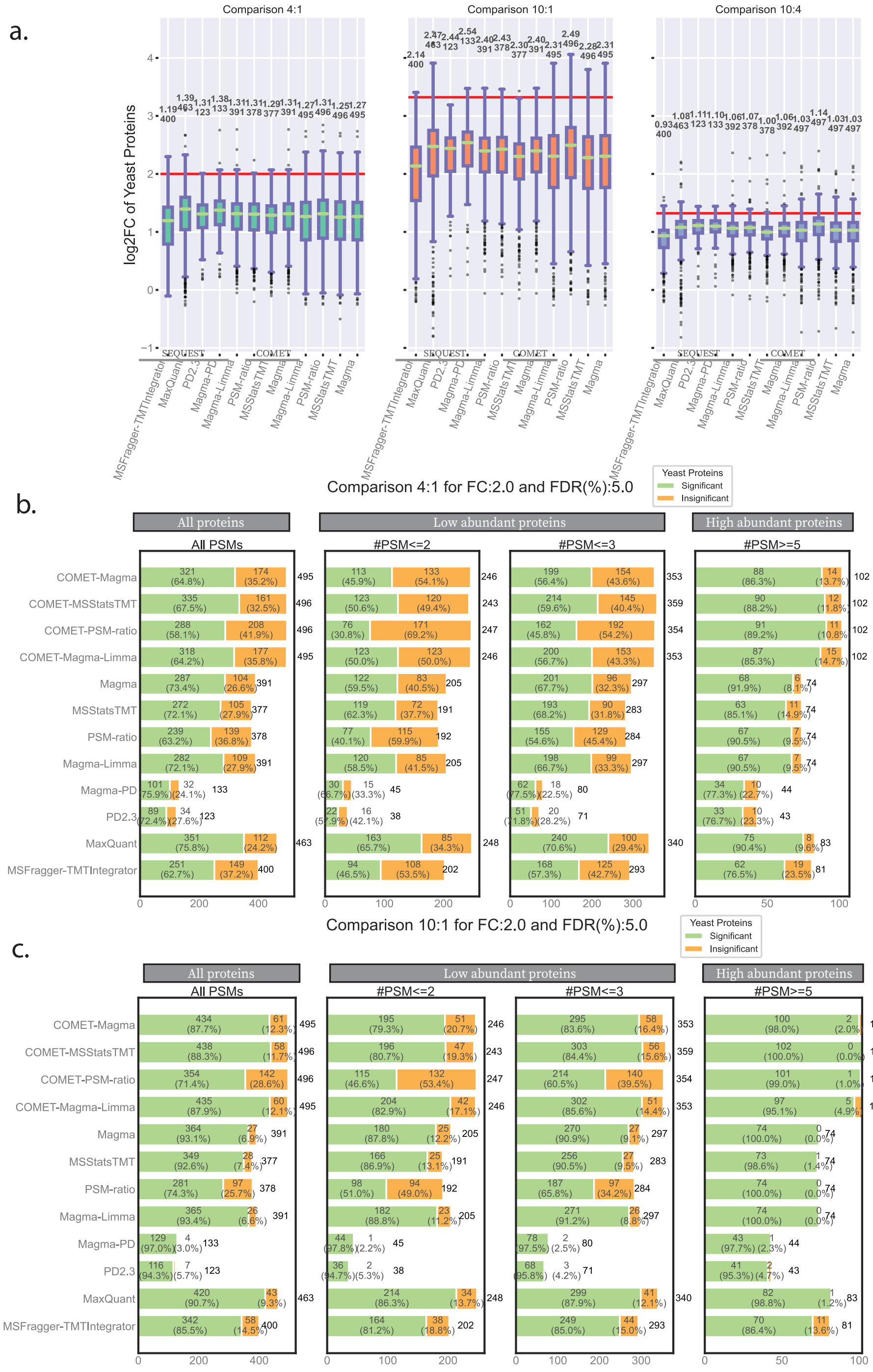


Supplementary Figure 3: **Performance evaluation of open-source and proprietary software(s) with MAGMa on the published phosphor-proteome benchmarking dataset.** (a) Distribution of log-2 transformed FC values for yeast proteins quantified with different combination of search and subsequent differential expression analysis tools. The three panels represent the three comparisons made (4:1, 10:1 and 10:4). The red line in each panel indicates the expected FC for that comparison. (b) Performance evaluation of various tools specifically on yeast proteins. It focuses on the tools' ability to detect an expected fold change (FC) of 4, comparing results across both low and highly abundant proteins. For each bar, in green are the number and proportion of yeast proteins that pass FC: 2 and FDR: 5% thresholds and in orange are number and proportion of yeast proteins that do not pass these thresholds. The number at the top of each bar is the total number of yeast proteins (c) Similar performance evaluation as (b) on yeast proteins on a comparison with an expected fold change (FC) of 10, comparing results across both low and highly abundant proteins.

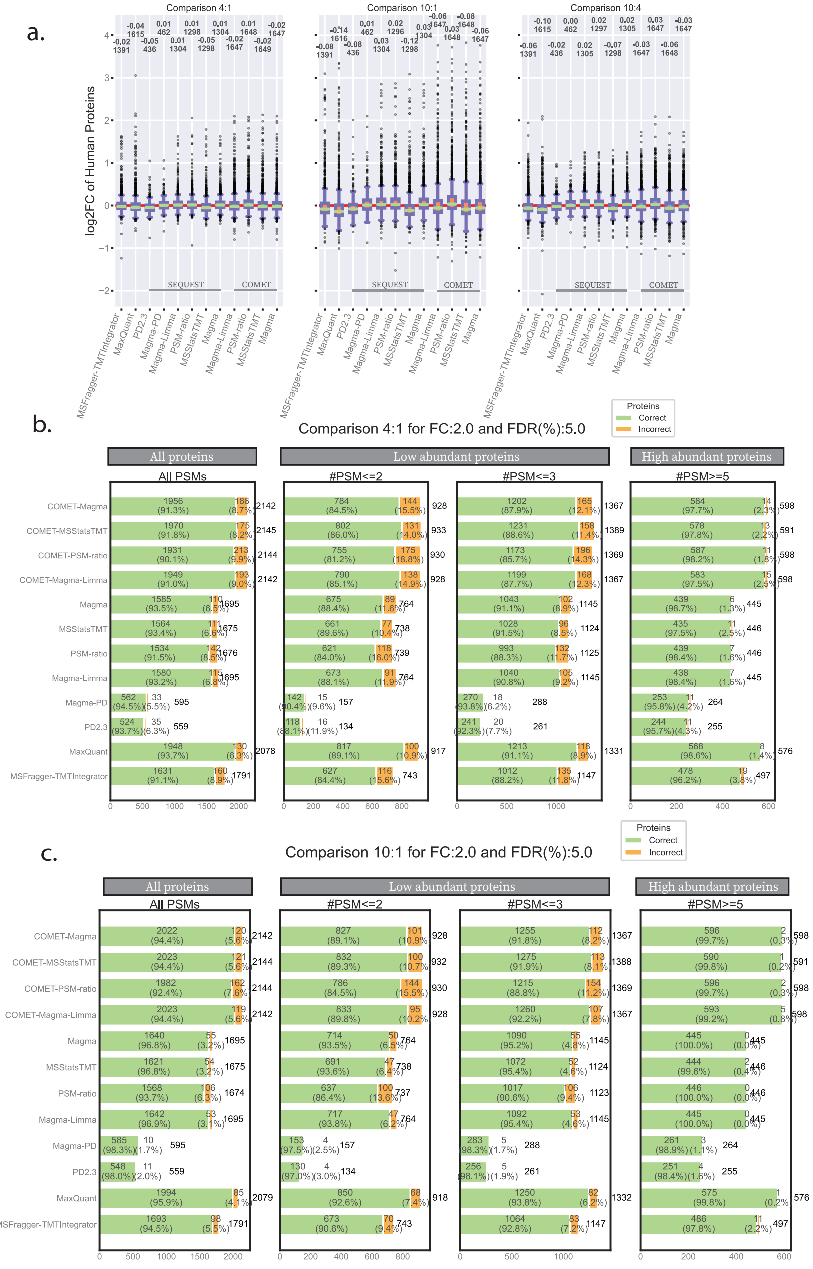


Supplementary Figure 4: **Performance evaluation of open-source and proprietary software(s) with MAGMa on the published phosphor-proteome benchmarking dataset.** (a) Distribution of log-2 transformed FC values for all human quantified with different combination of search and subsequent differential expression analysis tools. The three panels represent the three comparisons made (4:1, 10:1 and 10:4). The red line in each panel indicates the expected FC for that comparison. (b) Performance evaluation of tools on all proteins (yeast and human) on their ability to detect an expected fold change (FC) of 4 (yeast) and 1 (human), comparing results across both low and highly abundant proteins. For each bar, in green are the number and proportion of yeast proteins that pass FC: 2 and FDR: 5% thresholds and human proteins that do not pass these thresholds (correct calls). In orange are number and proportion of yeast proteins that do not pass these thresholds and number of human proteins that pass these thresholds (incorrect calls). The number at the top of each bar is the total number of proteins. (c) Similar performance evaluation as (b) on yeast and human proteins on a comparison with an expected fold change (FC) of 10 (yeast) and 1(human), comparing results across both low and highly abundant proteins.

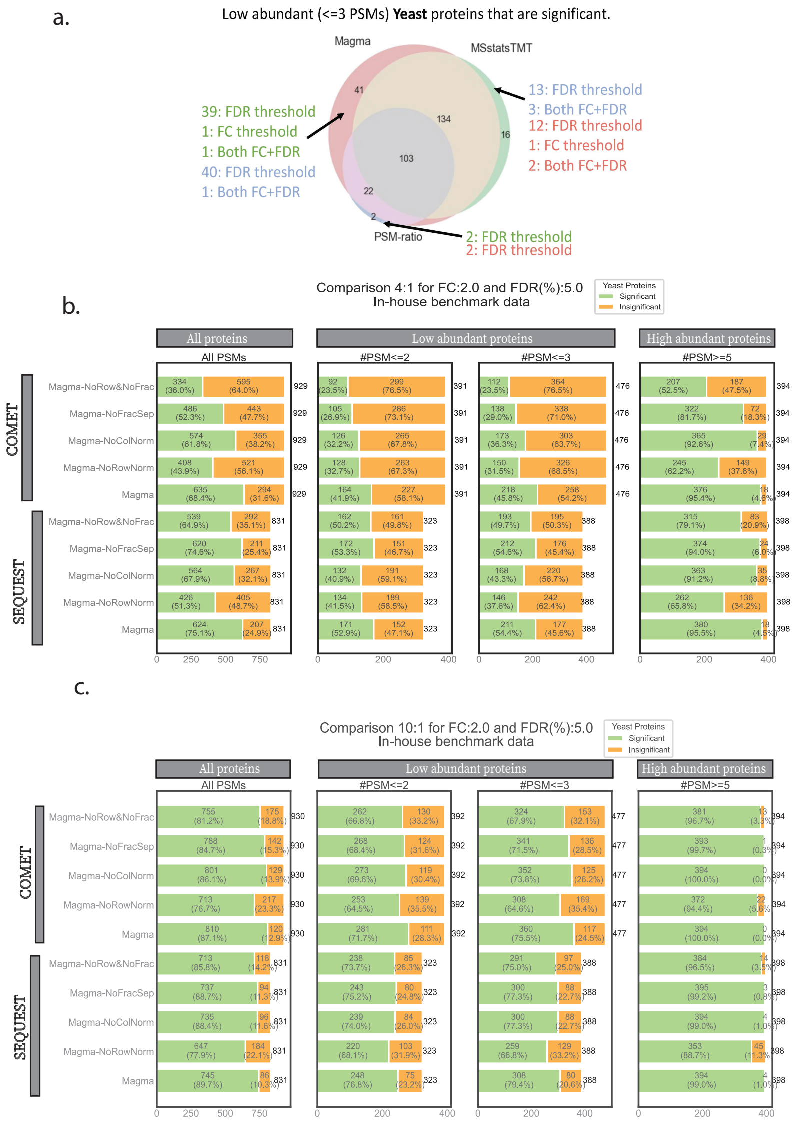


Supplementary Figure 5: **Ablation study on in-house benchmarking dataset with different MAGMa specifications.** (a) Venn diagram of lowly abundant proteins (with number of PSMs <= 3) quantified and identified as significantly enriched with MAGMa, MSstatsTMT, PSM-based methods using SEQUEST search. Threshold for calling significantly enriched is FC:1.5, FDR: 5%. Analysis of reasons for proteins to be missed out by different tools (green: MSstatsTMT, red: MAGMa, blue: PSM-based). These can either be due to FC not passing threshold, FDR not passing threshold, or both. (b) Similar to Figure 4 (a), ablation study done on yeast proteins for comparison with expected FC of 4. Thresholds of FC: 2 and FDR: 5% used. (c) Similar to Figure 4 (a), ablation study done on yeast proteins for comparison with expected FC of 10. Thresholds of FC: 2 and FDR: 5% used.

a.

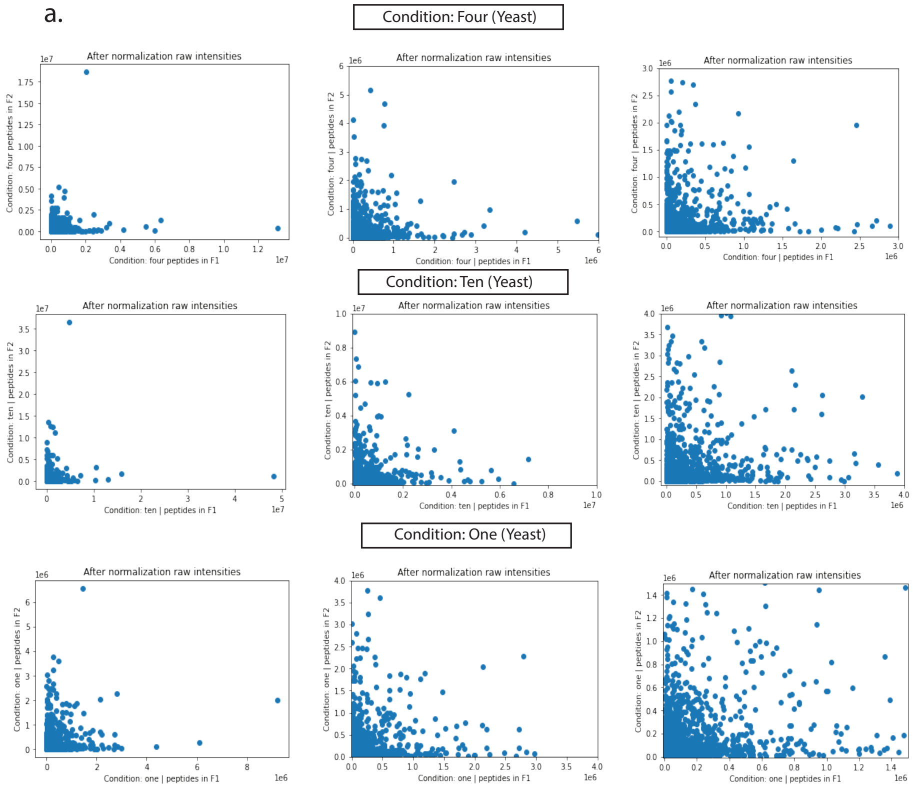


b.

| | Condition (Yeast, Human) | Pearson |
|---|---|---|
| Without row normalization | **1,1** | 0.261 |
| | **4,1** | 0.175 |
| | **10,1** | 0.147 |
| | **Condition (Yeast, Human)** | **Pearson** |
| Without row normalization , no column normalization | **1,1** | 0.261 |
| | **4,1** | 0.175 |
| | **10,1** | 0.147 |

Supplementary Figure 6: **Investigation of assumption that peptide measurements separated across fractions can be treated as independent.** (a) Scatter plot of all peptides identified in multiple fractions (randomly picked two if more), and their raw measurements (no normalization) taken from the same TMT channel (associated with the same reporter tag). This is done for all three conditions (B:4, A:10, or C:1) top to bottom and zoomed in from left to right. Top to bottom plots correspond to 3398, 3454 and 3250 peptides respectively. (b) For unique peptides (~3500) that came across multiple fractions, looked at Pearson correlation of raw reporter ion intensities by randomly picking two fractions/peptide and the same channel in both fractions.

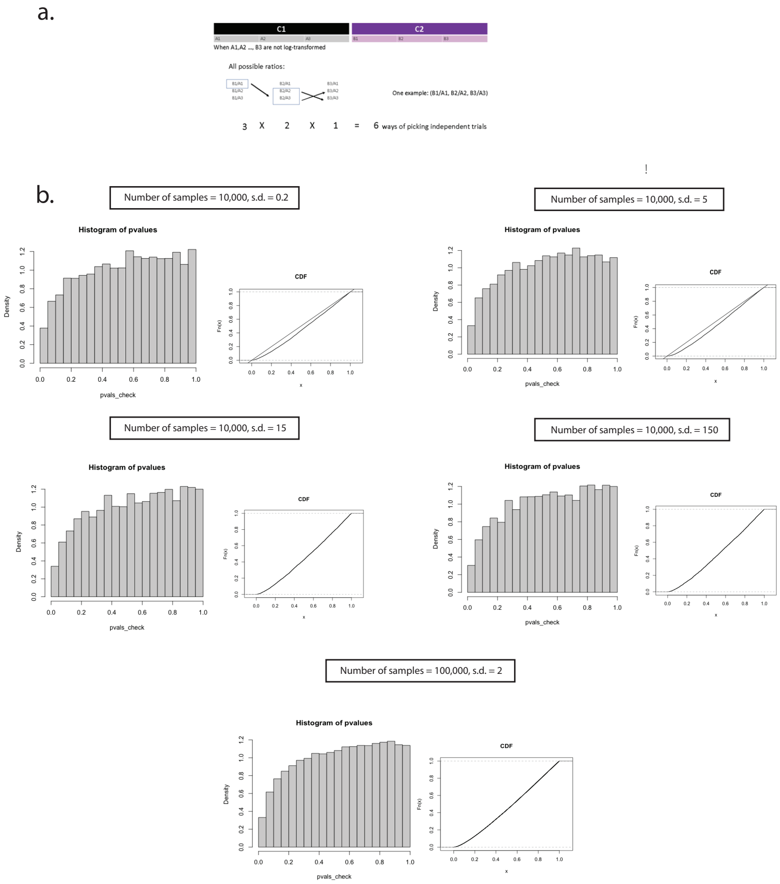


Supplementary Figure 7: **Investigation of feasibility of MAGMa-Limma approach of averaging *P*-values.** (a) Schematic explaining the number of ways in which independent trials per protein can be constructed given that its measurements come from three replicates per condition and found in one fraction. (b) Simulation to test assumption that average of *P*-values follows a uniform distribution given that *P*-values under the null hypothesis should follow a uniform distribution. This was conducted with increasing sample size and varying standard deviation (for treatment values). For each combination of sample size and standard deviation, the associated histogram of the averaged *P*-values (from three replicates per condition) and the cumulative distribution function is displayed.

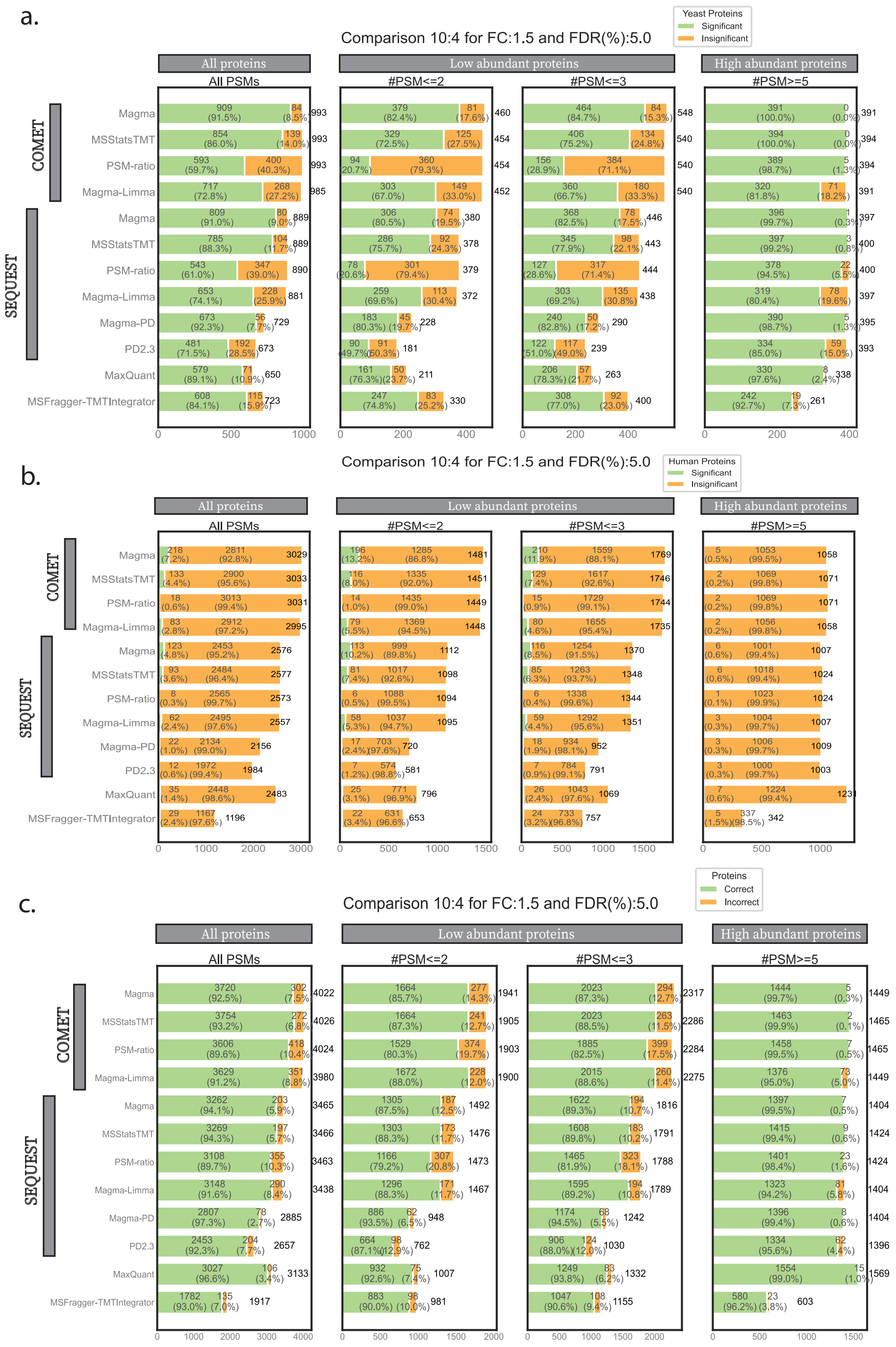


Supplementary Figure 8: **Impact of empirical bayes moderation on performance metrics on 10:4 comparison with threshold of FC: 1.5, FDR: 5%.** (a) Analysis on only yeast proteins as done in Figures before. (b) Same analysis as (a) but on human proteins only to showcase the larger number of false positive calls made as a result. (c) Same analysis done in previous figures as well as (a) and (b) but using all proteins.
